# Microtubule decay is a driver of neuronal ageing and a promising target for intervention

**DOI:** 10.1101/2023.01.11.523590

**Authors:** Pilar Okenve-Ramos, Rory Gosling, Monika Chojnowska-Monga, Kriti Gupta, Samuel Shields, Natalia Sanchez-Soriano

## Abstract

Natural ageing is accompanied by a decline in motor, sensory and cognitive functions, all impacting life quality. Ageing is the predominant risk factor for many neurodegenerative diseases, including Parkinson’s and Alzheimer’s disease. We need therefore to gain a better understanding of the cellular and physiological processes underlying age-related neuronal decay. However, gaining this understanding is a slow process due to the long time required to age mammalian or vertebrate model animals.

Here we introduce a new cellular model within the *Drosophila* brain where neurons show typical ageing hallmarks known from the primate brain, including axonal swellings, cytoskeletal decay, a reduction in axonal calibre and morphological changes arising at synaptic terminals. In the fly brain, these changes occur within just a few weeks, ideal to study the underlying mechanisms. We observe that decay of the neuronal microtubule cytoskeleton clearly precedes other ageing hallmarks. We show that the microtubule-binding factors Tau, EB1 and Shot, are necessary for microtubule maintenance in axons and synapses. Their functional loss during ageing triggers microtubule bundle decay followed by the decline in axons and synapses. Genetic manipulations that improve microtubule networks, slow down other neuronal ageing hallmarks and confer aged specimens with the ability to outperform age-matched controls. Our work suggests therefore that microtubule networks are a key lesion site in ageing neurons and offer promising opportunities to improve neuronal decay in advanced age.

## Introduction

Normal physiological ageing conveys a decline in sensory, motor and cognitive functions, with deterioration accelerating in the last decade of life. For example, short-term memory shows a slow deterioration with increasing age, which accelerates dramatically after the age of 70. Similarly, weakening of sensory functions, such as hearing and vision, become widespread amongst individuals over 80 years of age (Hedden and Gabrieli, 2004, Swenor et al., 2013, Gadkaree et al., 2016, Humes et al., 2013, Murman, 2015). The deterioration of these functions is characterised by behavioural changes and deficits. For example, the loss of sensory input is linked to a decline in social and physical activities leading to progressive social isolation, affecting daily living and individuals’ quality of life (Strawbridge et al., 2000).

The key factors which cause an age-related decline in brain function and cognitive behaviour are not well understood. However, a picture is emerging that suggests this is not due to widespread cell death (Walløe et al., 2014, Pakkenberg and Gundersen, 1997). Instead, distinct age-related alterations of neuronal structures, such as synapses, axons and dendrites, are proposed to be responsible for the widespread volumetric loss of brain tissue found in humans during ageing (Sherwood et al., 2011, Hedden and Gabrieli, 2004). Examples of such alterations in the mammalian brain include the progressive occurrence of atrophic axons, typically characterised by irregular membrane profiles and swellings (Fiala et al., 2007). Peripheral mouse motor axons display a marked decrease in axonal diameter with age (Ceballos-Baumann et al., 1999, Verdú et al., 2000, Borzuola et al., 2020). Additionally, in axons comprising the murine retina, varicosities and reduced synaptic content are observed upon age (Samuel et al., 2011). Synaptic structures also display morphological changes during ageing. For instance, in the rat hippocampus and mouse cerebellum, synapses develop discrete membrane swellings or oedemas, and exhibit a decrease in active zones (Rybka et al., 2019, Burke and Barnes, 2006, Fan et al., 2018). Understanding the causes for these structural changes will be pivotal in order to ameliorate the time-dependent decline of the nervous system.

A promising starting point is the microtubule (MT) cytoskeleton as an essential component of neurons, in particular their axons. MTs are arranged into parallel bundles that run uninterrupted all along axons; they form the highways for life-sustaining transport required for virtually all cell biological processes in axons, and they are essential elements driving morphogenetic processes (Kevenaar and Hoogenraad, 2015, Yogev et al., 2016, Prokop, 2020, Datar et al., 2019, Hahn et al., 2019). However, maintaining MT bundles long-term is a demanding task given the continued mechanical challenges imposed on them (Triclin et al.,2021, Atherton et al., 2018, Prokop, 2021, Dumont et al., 2015). Accordingly, MT density and organisation are reported to be altered in ageing axons of humans and other primates (Cash et al., 2003, Fiala et al., 2007).

Considering the unique dependency of cellular functions on MTs, we asked whether MT deterioration may be a cause for axon decay rather than a secondary consequence. *In vivo*, the regulation of MTs requires the function of MT-binding proteins (MTBPs) which regulate MT nucleation, polymerisation, disassembly, stabilisation and cross-linkage, posttranslational modification and repair (Kapitein and Hoogenraad, 2015, Voelzmann et al., 2016a, Hahn et al., 2019, Gasic and Mitchison, 2019). Loss of balance in this complex regulatory network of MTBPs could explain MT decay and neuronal atrophy reported during ageing (Cash et al., 2003, Brunden et al., 2017, Kounakis and Tavernarakis, 2019a, Czaniecki et al., 2019, Prokop, 2021). We propose that gaining a better understanding of MT regulation and its roles during axonal maintenance and decay provides a promising path to gain new insights into age-related neuronal decay and unveil new therapeutic strategies.

To be able to deal with the complexity of axonal cell biology and of MT regulatory mechanisms, we used the model organism *Drosophila melanogaster.* Importantly, *Drosophila* shows clear signs of severe ageing within a matter of few weeks, including diminished stress resistance, changes in behaviour, altered metabolism, reduced barrier function in the gut, and compromised cardiac function (Piper and Partridge, 2018). Ageing has also been reported to impact *Drosophila* neurons, including age-related changes in mitochondrial dynamics, synaptic content and diminished neuronal function (Hussain et al., 2018, Vagnoni and Bullock, 2018, Vagnoni et al., 2016, Bhukel et al., 2019, Banerjee et al., 2021, Rhodenizer et al., 2008, Birnbaum et al., 2020).

Using this model, we performed a detailed characterisation of neuronal ageing in cells situated within the fly visual system. We found prominent ageing hallmarks remarkably similar to those observed in mammals, including axonal swellings, altered axon diameters, cytoskeletal and synaptic decay. The earliest symptom we observed was MT bundle decay (discontinuity, disturbances in the bundle organisation and loss of MT volume) which correlated with decreased functions of several MTBPs. Genetic reduction of these MT regulators induced ageing-like MT defects and exacerbated age-dependent neuronal decay, whereas genetic manipulations stabilising and enhancing MT networks protected neurons from developing hallmarks of ageing, thus demonstrating a causative link between MT deregulation and neuronal decay.

## Results

### Age alters the morphology of axons and synapses

To understand how axons and synaptic terminals are changing with age and what drives this process, we set out to establish an *in vivo* cellular model which would allow us to study the cell biology of ageing. We focused on the *Drosophila* visual system, in particular on T1 neurons which are second order interneurons in the medulla rind (Fig. 1A). T1 neurons form projections that bifurcate into one branch arborising in layer 2 of the medulla where they contact several synaptic partners, and another projecting into the lamina (Myers Gschweng et al., 2019, Takemura et al., 2015) (Fig. 1A). Several properties make the medulla-localised axons of T1 neurons ideal for subcellular studies: they are large calibre axons with characteristic synaptic terminals in stereotyped medulla locations; they are easy to image and manipulate due to their proximity to the surface of the tissue; and numerous genetic tools are available to facilitate their effective visualisation and manipulation.

**Figure 1.**
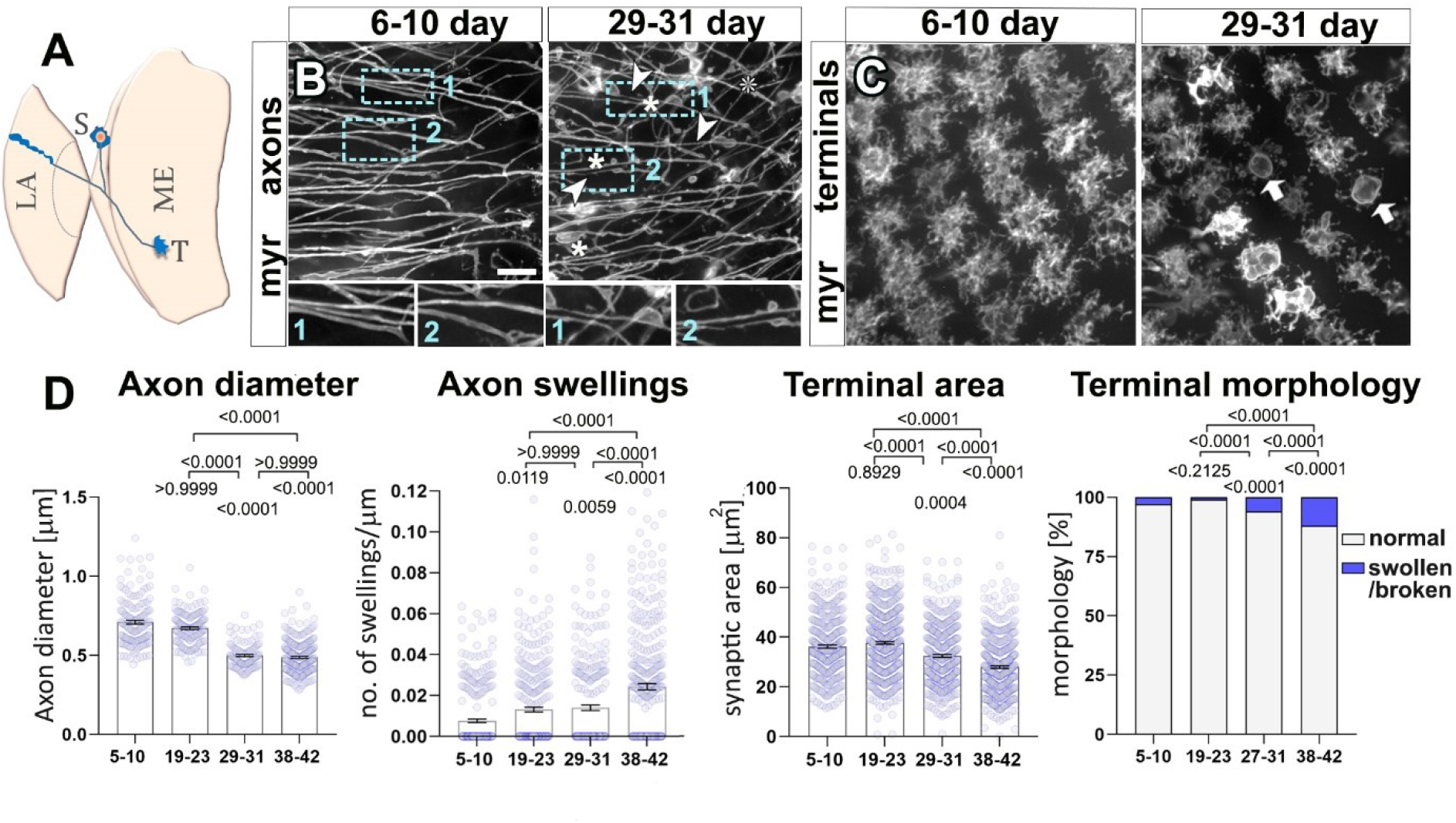
Axons and synaptic terminals within the *Drosophila* visual system deteriorate during ageing. (**A**) Representation of an optic lobe region of the adult *Drosophila* brain depicting a soma (S) of a T1 neuron which has a projection that bifurcates terminating in the medulla (ME; T, terminals analysed in this study) and in the lamina (LA). (**B,C**) Axons (B) and terminals (C) of the T1 medulla projections from young (6-10 day) and old specimens (29-31 day) labelled with the plasma membrane marker myr-Tom (myr). In aged specimens, axons show thinning (arrow heads) and swellings (asterisks; dashed blue boxes shown as 1.7-fold magnified images below), and axonal terminals appear broken down and with swellings (arrows) and overall reduced area. (**D**) Quantifications of phenotypes shown in B and C, with age groups in days after eclosure, indicated on the X-axes. In the left three graphs, data points are shown in lilac and as mean ± SEM. P-values obtained via Kruskall-Wallis ANOVA multiple comparison tests are indicated in the graphs. For terminal morphology, data are represented as distribution of normal versus swollen/broken synapses; significance obtained via Chi-square test is indicated above. Data were taken from a minimum of 10 specimens, 100 axons or 300 terminals per age group. For detailed statistical values and genotypes see table S1. Scale bar in B represents 10 μm in B, and 14 μm in C.

To study the medulla axons of T1 neurons, we labelled the membrane via targeted expression of the fluorescently tagged membrane marker myristoylated-Tomato (myr-Tom) driven by the *GMR31F10-Gal4* line (Qu et al., 2019). Labelled axons were visualised via an *ex vivo* live imaging method (see Materials and Methods). We performed comparative analyses between specimens of different ages. Since the range of temperatures at which *Drosophila* can be reared is variable and higher temperatures facilitate faster specimen ageing, we opted to age *Drosophila* at 29°C, a temperature that is ecologically relevant and allows faster experimentation (Soto-Yéber et al., 2019). Under these conditions, we found that T1 axons become thinner (measured as a decrease in their diameter; see Materials and Methods), their originally smooth membranes become irregular, and they develop frequent axonal swellings (Fig. 1B). These changes were widely abundant in flies aged over 29 days after eclosure (DAE, Fig. 1D). Other neurons within the visual system also undergo similar age-related changes in axons, as shown in L2 lamina neurons (Fig. S1), suggesting that these alterations are not T1-dependent and may be widespread across other *Drosophila* neuronal cell-types.

In addition to morphological changes in the shaft of axons, we found that ageing altered the morphology of axonal terminals containing synapses. In young specimens, T1 terminals in layer 2 of the medulla neuropile consist of hand-shaped, dense arborisations (Fig. 1A and C). In old specimens (starting at ~30 days old), these arborisations become irregular and less compact, resulting in an overall decrease in synaptic area and, in extreme cases, apparent fragmentation of the synaptic membranes (Fig. 1C and D). Additionally, synaptic terminals appear swollen, adopting a highly spherical and varicose shape (Fig. 1C and D). These changes occur in the absence of neuronal death, as suggested by quantification of T1 neuron nuclei labelled with RedStinger (Bischoff and Cseresnyés, 2009) (Fig. S2). These ageing alterations occurring in *Drosophila* show similarities to transformations which occur in the murine retina, motor axons, and the primate brain, which all display axonal varicosities and reduced axonal terminals with age (Samuel et al., 2011, Ceballos-Baumann et al., 1999, Verdú et al., 2000, Borzuola et al., 2020, Fiala et al., 2007).

**Figure 2.**
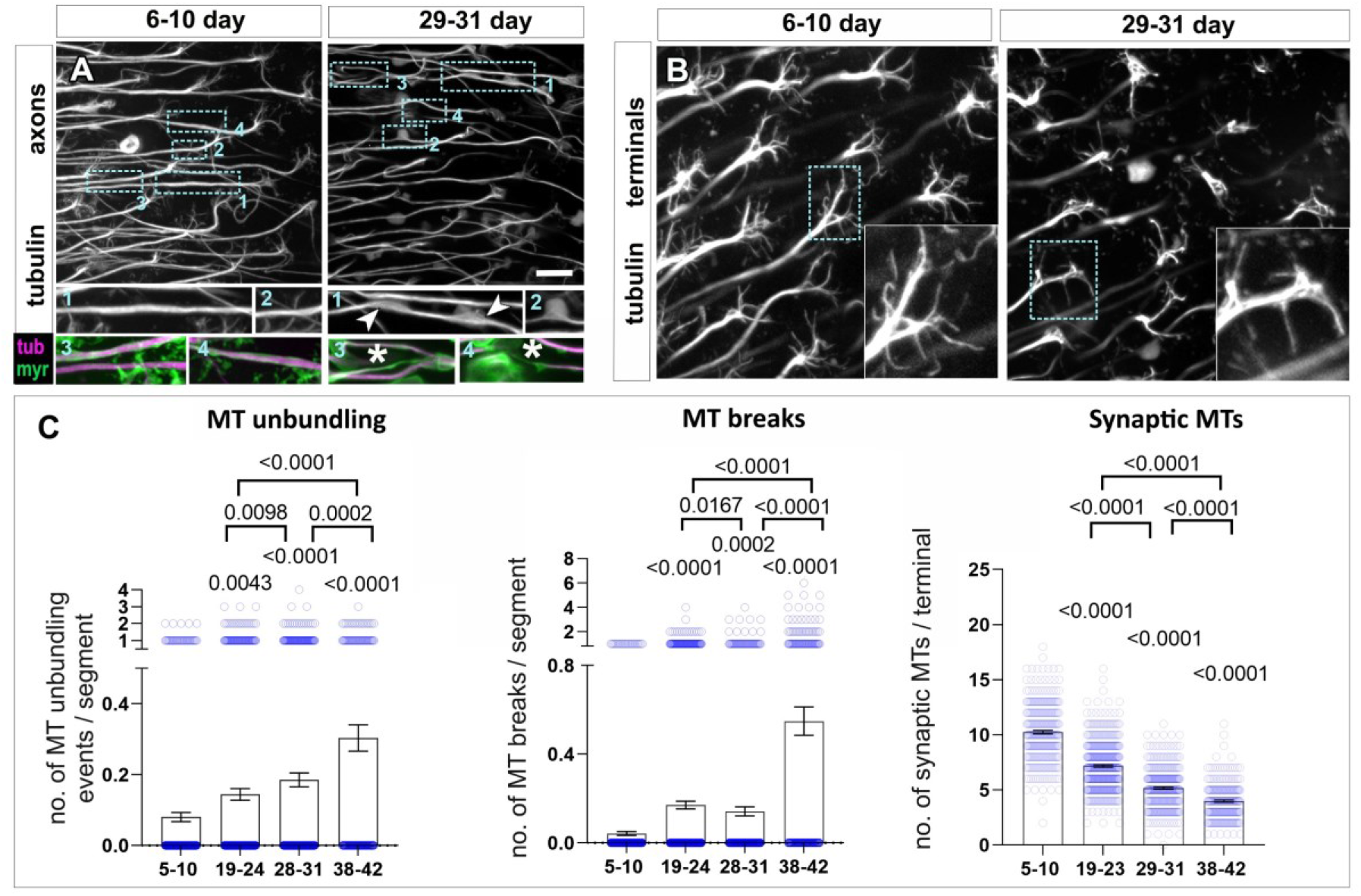
MT aberration precedes axonal and synaptic decay during ageing. (**A-B**) T1 neuron axons (A) and synaptic terminals (B) in the medulla region of flies which were either young (6-10d of adult life) or old (29-31d of adult life) and labelled with GFP-tagged α-tubulin (tubulin or tub., in grey, and magenta in the insets) and the plasma membrane marker myr-Tom (myr, green). In A, cyan encircled boxes are shown 2.3-fold magnified below with old flies showing MT unbundling (insets 1 and 2 in A) as well as breaks and thinning of MTs (insets 3 and 4 in A) compared to young individuals. In B, old flies also show a reduction of splayed MTs in synaptic terminals (boxed areas shown as 2-fold magnified insets). (**C**) Quantifications of phenotypes shown in A and B, with age groups indicated below X-axes; data points are shown as blue circles and as mean bars ± SEM; p-values obtained via Kruskall-Wallis ANOVA multiple comparison tests are indicated above. Data were taken from a minimum of 17 specimens per age group. For detailed statistical values see table S4. Scale bar in A represents 10 μm in A and 25 μm in B.

To demonstrate that these alterations were not a consequence of long-term expression of the membrane marker myr-Tom, we used the conditional expression system Gal4/Gal80^ts^ (Pfeiffer et al., 2010, Suster et al., 2004) (see Materials and Methods) which enabled us to restrict expression of the membrane marker to equal short time periods in both young and old flies. This system relies on the ubiquitous expression of the transcriptional repressor Gal80, which contains a temperature-sensitive mutation that renders it inactive at 29°C. Thus, at 29°C, Gal80 is inactivated, allowing Gal4 to freely induce transcription of UAS-dependent transgenes, whereas in flies reared at 18°C, expression of the transgene is repressed by active Gal80. To this end, we aged flies for either 57 to 60 or 4 to 7 days at 18°C in the absence of myr-Tom (marker expression is suppressed at 18°C), then incubated both young and old flies at 29°C for 4 days to enable short-term expression of myr-Tom, before dissecting and imaging the brains (Fig. S3). We found that axons from old specimens displayed: reduced diameters, irregular membrane profiles, increased swellings and altered synaptic morphology, when compared to young ones. This indicated that the defects observed using this model were a consequence of ageing and not of long-term stress due to continuous prolonged myr-Tom expression. These results also show that significant ageing alterations found in axonal and synaptic terminals are independent of the temperature *Drosophila* is reared at.

Taken together, our data demonstrated that significant structural changes occur in both axonal and synaptic compartments during ageing. Aged neurons display a decrease in the diameter of axons which develop swellings in addition to alterations in synaptic morphology, suggesting that axonal and synaptic integrity are compromised.

### Changes in the MT cytoskeleton precede axonal and synaptic decay during ageing

The MT cytoskeleton is a key determinant of cell shape and of the morphology of axons and synaptic terminals (Datar et al., 2019, Fan et al., 2017, Saunders et al., 2022). MTs are also suggested to be key factors in regulating the calibre of axons, especially in conditions of low abundance or absence of neurofilaments (Stephan et al., 2015, Perge et al., 2012, Friede and Samorajski, 1970, Prokop, 2020). We therefore hypothesised that MTs can play a key role in the context of age-related neuronal defects observed with our model. To this end, we first investigated whether ageing acutely affects the MT cytoskeleton of neurons.

To visualise axonal and synaptic MTs in T1 neurons, we expressed GFP-tagged α-tubulin (tubulin::GFP, Grieder et al., 2000). As anticipated, axonal MTs of T1 neurons in young adults displayed a prominent, continuous and uniform bundle along the axon, which splays into multiple thinner bundle branches at the synaptic terminal (Fig. 2A and B). However, in older specimens, we observed that axonal MTs lose their tight, parallel arrangements, deviating from their bundles leading to disorganised foci (MT unbundling). Moreover, there were regions within axons which appeared to contain thinner MT bundles and, in extreme examples, devoid of MTs (MT breaks) (Fig. 2A and C). We confirmed the age-dependent overall thinning of axonal MT bundles using high-end confocal microscopy as well as stimulated emission depletion super-resolution microscopy (STED). By measuring the cross-section of MT bundles within T1 axons (as an indication of the bundle diameters), we found a significant decrease in old versus young specimens when imaged with both techniques (Fig. S4). In parallel, we also observed that the splayed synaptic MTs often appeared fragmented and decreased in number with age (over 50% reduction from 38 days; Fig. 2B and C). Upon assessment of the ageing phenotypes at multiple timepoints, it was revealed that axonal and synaptic MT bundle deterioration is already prominent from 19 days onwards, demonstrating that MT defects occur prior to the onset of morphological changes, such as thinning of axons, increased swellings and synaptic alterations (detectable not before about 30 days; Fig. 2C vs Fig. 1D).

In order to discard any confounding contributions to such phenotypes due to long term expression of tubulin::GFP, we utilised the same approach as carried out for myrTom: the Gal4/Gal80^ts^ system was used to control temporal expression of tubulin-GFP for an equal amount of days in young and old specimens. Under these conditions of time-controlled expression, aged specimens still developed the same age-related MT phenotypes (Fig. S5), thus ruling out long-term expression of tubulin-GFP as the cause of the described ageing phenotypes.

To validate whether these various observed phenotypes are the consequence of ageing, we knocked down *methuselah* (*mth*), the loss of function of which is known to prolong lifespan by influencing the dTOR pathway (Lin et al., 1998). Upon loss of *mth* in our model, all measured ageing hallmarks found in old animals were significantly improved, including swellings and axonal thinning as well as MT deterioration in the axons and synaptic terminals; Fig. 3).

**Figure 3:**
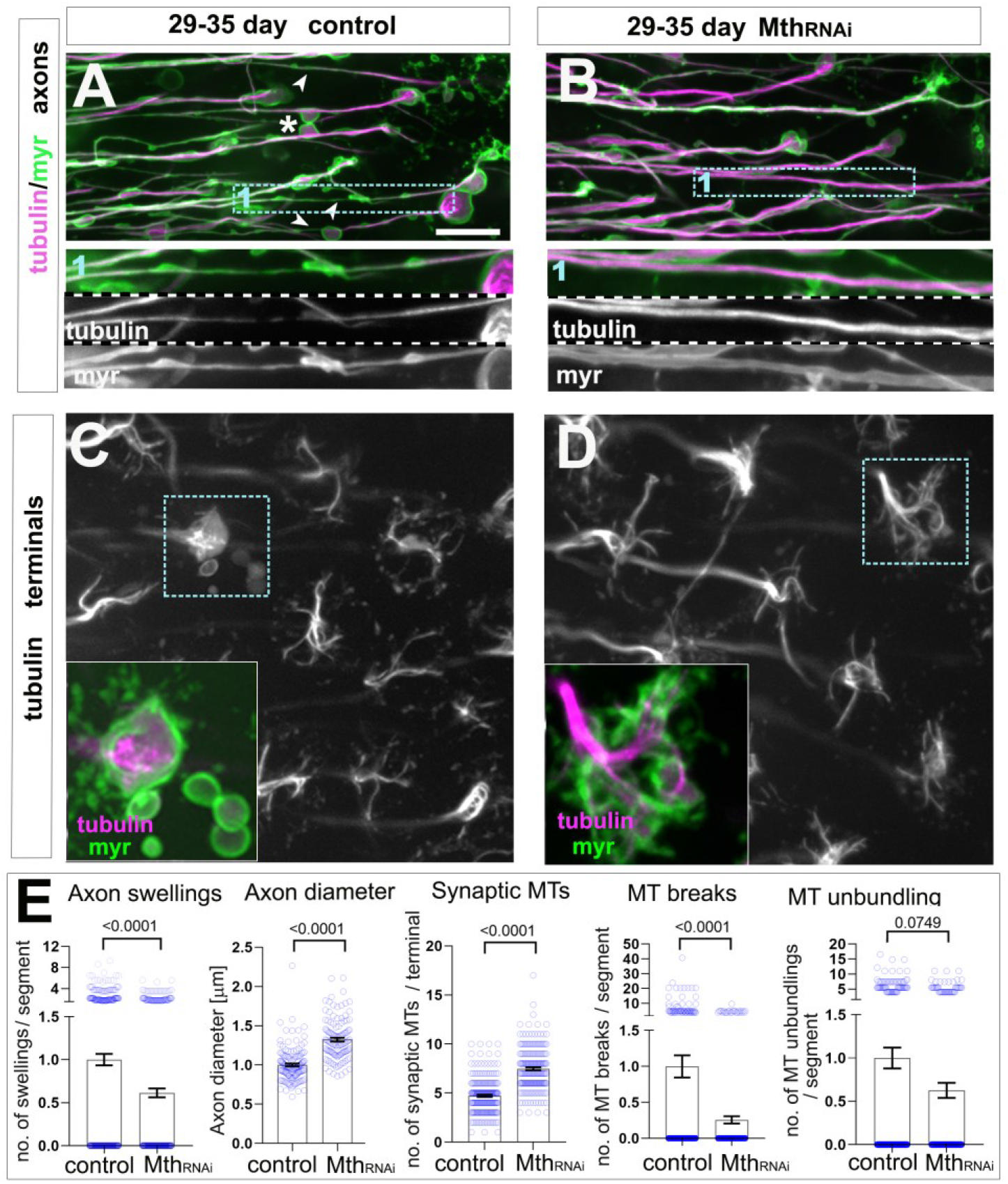
Knockdown of the ageing gene *mth* improves neuronal ageing hallmarks and MT decay. (**A-D**) T1 axons (top) and synaptic terminals (bottom) in the medulla of old specimens of 29-35 days, are labelled with GFP-tagged α-tubulin (tubulin, magenta in A and B and greyscale in C and D) and the plasma membrane marker myr-Tom (myr, green in A and B). Ageing phenotypes in old specimens in A include axon swellings (asterisks), axon thinning (arrow heads) and MT breaks/thinning (boxed area 1 shown as 2-fold magnified inset); as well as reduction of splayed MTs in synaptic terminals shown in C (boxed areas shown as 2-fold magnified insets). T1 specific knockdown of the ageing gene *mth* (Mth^RNAi^) in B and D, suppresses phenotypes in ageing neurons in aged-matched specimens. (**E**) Quantifications of phenotypes shown in A-D, in the absence or presence of Mth^RNAi^ indicated below X-axes; data points are shown as blue circles and as mean ± SEM; p-values obtained via Mann-Whitney test are indicated above. Data were taken from a minimum of 16 specimens per group (except for axonal diameter and synaptic MTs where a minimum of 10 specimens were used). For detailed statistical values see table S6. Scale bar in A represents 10 μm in A and B, and 25 μm in C and D.

Our results indicate that the hallmarks reported here are the consequence of physiological ageing. Furthermore, MT bundle deterioration occurs before other subsequent structural changes, such as the onset of swelling and axonal thinning, suggesting that MT bundle deterioration is an important event implicated in axon and synaptic decay observed during ageing.

### The turnover of axonal MTs decreases with age

The thinning of axonal MT bundles and the decrease in synaptic MTs we observed in old specimens, may suggest that MT turnover is compromised with age. Mechanisms of MT nucleation, polymerisation, stabilisation and bundling within axons have been the subject of multiple studies conducted primarily in developing axons (Kapitein and Hoogenraad, 2015, Voelzmann et al., 2016a, Hahn et al., 2019, Gasic and Mitchison, 2019). However, to our knowledge, very little is known about MT turnover in mature axons which can be expected to be an important mechanism to counteract wear and tear, imposed for example by motor proteins (see Introduction).

To investigate MT turnover in mature neurons, we performed a pulse-chase experiment in which tubulin::GFP was expressed during a restricted period of 4 days in either young or old specimens, after which we examined to what degree *GMR31F10-Gal4-driven* tubulin::GFP was incorporated into the axonal MT bundle. To achieve this, we utilised the conditional Gal4/Gal80^ts^ expression system: flies were maintained at 18°C (tubulin::GFP expression is inactive) for either 0 to 3 (young specimens) or 53 to 56 DAE (old/aged specimens), followed by 4 days at 29°C (tubulin::GFP expression is active). In young and old specimens, tubulin::GFP was clearly incorporated within axonal MTs, indicating that MTs continue to turnover in adult T1 neurons at both timepoints (Fig. 4A-B). To assess the level of tubulin::GFP incorporation, we quantified the intensity of GFP in axonal MT bundles, and to exclude potential artefacts due to changes in driver activity, we normalised tubulin::GFP to co-expressed myr-Tom. These analyses clearly revealed a significant decrease in GFP intensity in axons from old flies when compared to the younger ones (Fig. 4C).

**Figure 4:**
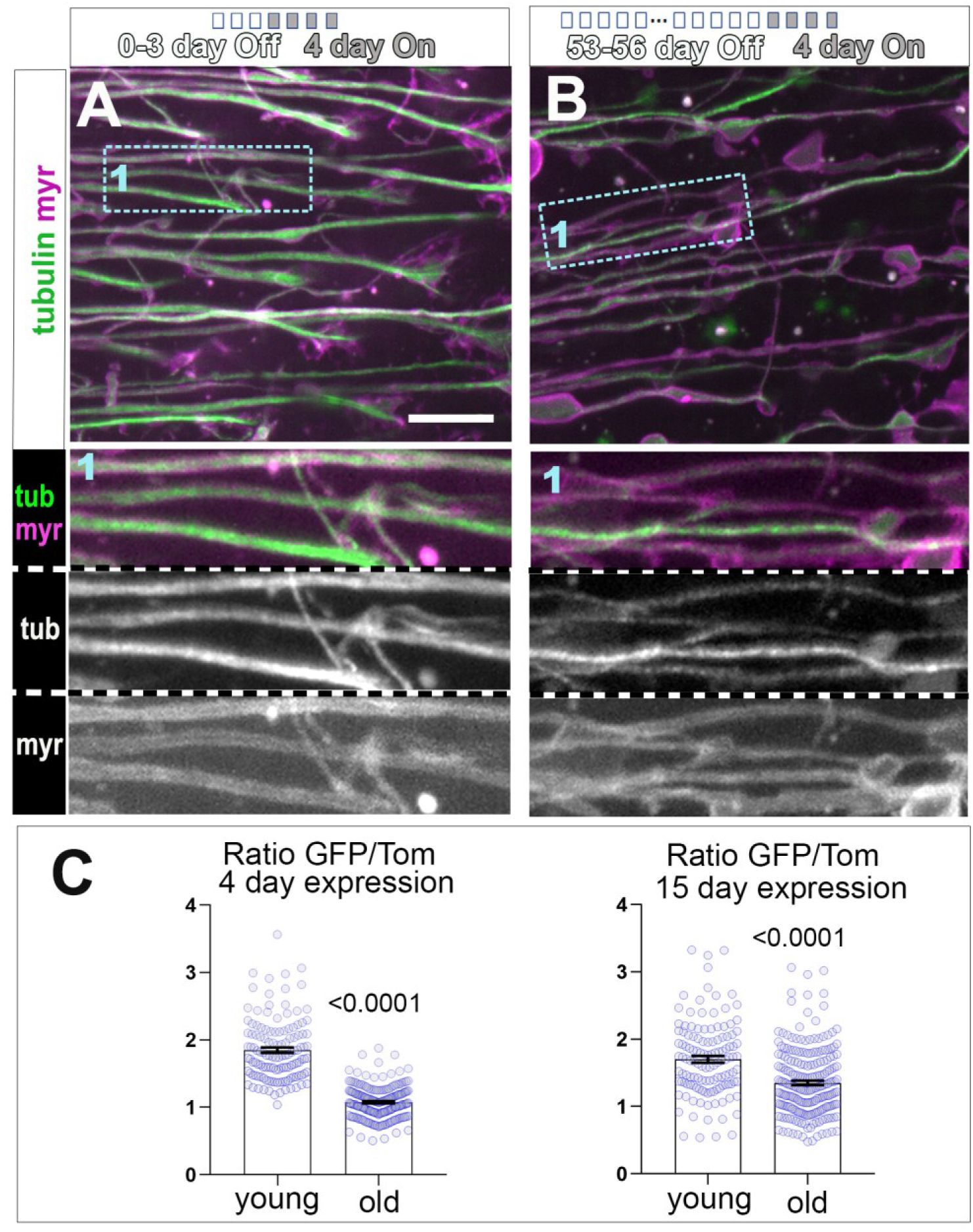
Decreased presence of tubulin-GFP at axonal microtubules in a pulse chase experiment suggests changes in MT turnover with age. (**A-B**) T1 axons in the medulla of flies expressing GFP-tagged α-tubulin (tubulin, in green) and the plasma membrane marker myr-Tom (myr, magenta), using the UAS/Gal4/Gal80^ts^ system. The induction of gene expression in the adult brains is initiated and restricted to 4 days by the shift of temperature from 18 to 29°C. For this, flies were kept at 18°C throughout development and adult life until the last 4 days before imaging, at which point they were shifted to 29°C to initiate gene expression. Young specimens (A; 4-7 day old flies with 0-3 days at 18°C “Off” + 4 days at 29°C “On”) are compared to old specimens (B; 57-60 day old flies with 53-56 days at 18°C “Off” + 4 day at 29°C “On”). The level of Tub-GFP incorporated within axons is diminished in old specimens (B compared to A and magnified blue boxes shown as 2-fold magnified insets). (**C**) Quantifications of the ratio between GFP-tub and myr-Tom (Ratio GFP/Tom) in two experimental conditions (under 4 days or 15 days expression regime), with young versus old indicated on the X-axes; Data points are shown as blue circles and as mean ± SEM; p-values obtained via Mann-Whitney test are indicated above. Individual data are taken from at least 5 or 6 specimens per age group for 4 or 15 day expression respectively. For detailed statistical values Table S7. Scale bar can be found in A bottom left (A-B 10 μm).

These data strongly suggest that MT turnover is diminished during ageing, which could explain the deterioration of the axonal MT bundle we observed in ageing flies.

### MT-binding activities of EB1 and Tau decrease with age

Next, we set out to investigate the mechanisms that could lead to the decrease in MT turnover with age. The maintenance of MT bundles is expected to require the function of MT-binding proteins, such as the MT lattice-binding factor Tau and the MT plus end-binding factor EB1. EB1 is a scaffold protein preferentially localising to polymerising MT plus ends where it mediates the binding of factors involved in polymerisation dynamics and the guidance of extending MTs into organised bundles (Alves-Silva et al., 2012, Hahn et al., 2021, van de Willige et al., 2016, Voelzmann et al., 2016a, Duellberg et al., 2016). Tau stabilises MTs along their lattice, and it promotes MT polymerisation by maintaining EB1 at extending MT plus ends (Voelzmann et al., 2016b, Hahn et al., 2021b, Iwata et al., 2019); the aberrant localisation of Tau is a hallmark of ageing in the human brain and correlates with memory dysfunction (Harrison et al., 2019, Reas, 2017).

We started by investigating age-dependent changes in EB1 and Tau localisation in the brain. We visualised EB1 performing antibody staining of whole brains. In the medulla of young specimens, we found that EB1 staining appeared as distinct speckles throughout, with regions of short, continued strokes within areas containing axons (Fig. 5A). Such patterns of expression likely represent EB1 bound to the plus end and/or discrete regions along MT lattices as observed upon MT repair (Hahn et al., 2021, Triclin et al., 2021). Notably, EB1 localisation to axons was weaker in brains dissected from old flies, as suggested by a reduction in EB1 intensity in axonal regions within the medulla (Fig. 5A and B).

**Figure 5:**
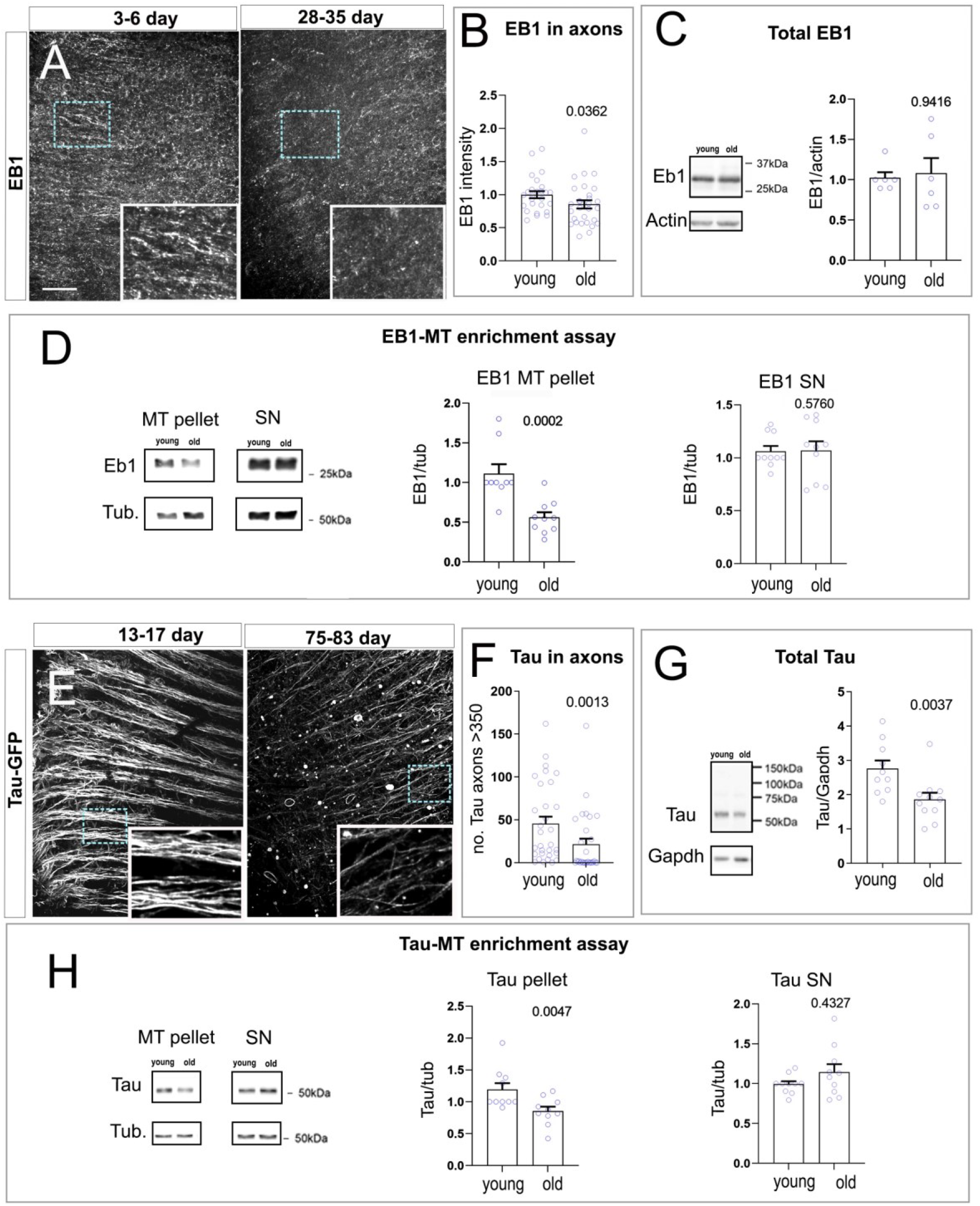
The function of EB1 and Tau is altered during ageing. (**A**) Axonal region of the medulla of young (3-6 day) and old (28-35) flies labelled with anti-EB1. (**E**) Axonal region of a medulla labelled with endogenous GFP-tagged Tau, at two different timepoints (young 13-17 and old 75-83 day old, both ager at 25°C). In aged specimens, axons show a decrease in EB1 and GFP-Tau (dashed cyan boxes in A and E are shown as 2 and 2.3-fold magnified insets respectively). Images as in A and E are used for (**B**) quantifications of EB1 signal intensity and (**F**) of the number of axons with high levels of GFP-Tau (above 350 units of intensity). Age categories as explained above are indicated below X-axes. Data points represent medullas and are shown as blue circles and as mean ± SEM; P-values obtained by Mann-Whitney test are indicated above. Data were taken from a minimum of 17 specimens per age group. (**C** and **G**) Total EB1 and Tau by western blot reveals a decrease in Tau during ageing, while the level of EB1 remains constant. Age categories are indicated below X-axes (4-9 and 29-32 for C and 8-10 and 29-31 for G). Data points represent independent lysates and are shown as blue circles and as mean ± SEM; P-values obtained by Mann-Whitney test are indicated above. (**D** and **H**) Microtubule-binding assays were performed using head extracts from young and old specimens, followed by fractionation to obtain a MT-enriched pellet and a soluble supernatant (SN). Both the MT and SN fraction were probed with anti-EB1, anti-Tau and anti-Tubulin. Data points represent independent microtubule-binding assays and are shown as blue circles and as mean ± SEM; P-values obtained by Mann-Whitney test are indicated above. Quantifications confirm that the amount of EB1 and Tau bound to MTs decreases with age. For detailed statistical values see table S8. Scale bar in A represents 20 μm in A and E.

Tau antibody staining proved to be unreliable in whole brains. As an alternative strategy, we used the viable *tau^wee304^* allele which contains a GFP inserted into its genomic locus (Stone et al., 2008). In young adult *tau^wee304/+^* specimens, Tau^wee304^ was predominantly localised to axons within the medulla, exhibiting a filament-like pattern, consistent with binding along MT lattices (Fig. 5E). However, this pattern changed in aged medullas which displayed a stark decrease in Tau^wee304^ GFP intensity in axons (Fig. 5 E and F). In addition, we observed an increase in aggregate-like structures (Fig. 5E). This relocalisation of *Drosophila* Tau shows striking similarities to alterations observed in aged human individuals or upon certain tauopathies including Alzheimer’s disease (Schöll et al., 2016, Harley et al., 2021)

Reduced MT association of EB1 and Tau in ageing neurons may reflect an overall decrease in their levels. We therefore quantified Tau and EB1 protein levels in extracts from *Drosophila* heads isolated at different ages. We found that the level of EB1 remained unaffected in old specimens (Fig. 5C), while there was a decrease of total Tau in old specimens (Fig. 5G). The decrease in Tau could be due to either decrease in expression or a sequestration of Tau in aggregates which may have not been detected in our western blot studies.

To further understand whether the MT regulatory function of Tau and EB1 may be altered during ageing, we compared the MT-binding properties of these factors in adult young and old brains. For this, we performed a standard MT binding spin-down assay using extracts from fly heads at different ages (Cowan et al., 2010). We found that the amount of EB1 and Tau detected in the MT-enriched fraction from old brains significantly decreased when compared to young counterparts (Fig. 5D and H), implying that there is less EB1 and Tau bound to MTs in old specimens. This suggests that the ability of EB1 to bind MTs decreases with age and is consistent with the reduced EB1 intensity we observed in aged axons, while the axonal Tau loss in aged brains may be due to both diminished MT binding and a global decrease in protein levels in the brain.

### Loss of MT regulators exacerbates ageing hallmarks

Our results might suggest that an age-related decrease in the function and/or abundance of key MT regulators could be the cause of MT loss and unbundling, and even of the subsequent axonal and synaptic decay that occurs in the ageing process. To assess this possibility, we performed targeted expression in T1 neurons of previously validated RNAi constructs against several essential MT regulators: Tau, EB1 as well as the *Drosophila* spectraplakin Short stop (Shot) as a further essential MT regulator (Voelzmann et al., 2016b, Alves-Silva et al., 2012). T1-specific knock-down of Tau in old specimens compared to age-matched controls, resulted in a significant increase in the number of sites where MT bundles were disturbed or displayed breaks. These MT phenotypes correlated with a rise in the number of axonal swellings (Fig. 6C).

**Figure 6:**
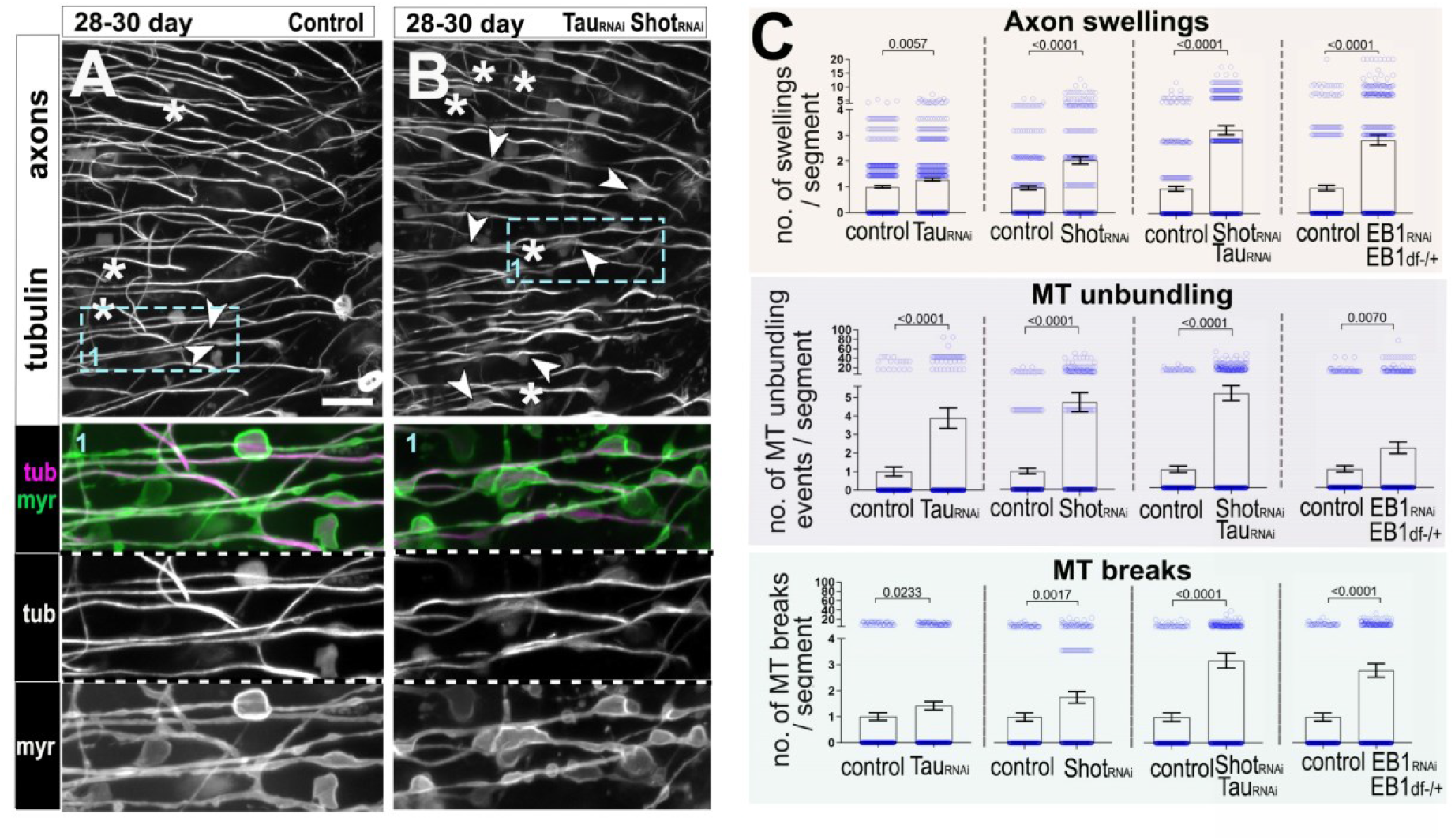
Reduction of Eb1, Tau and Shot exacerbates age-related MT decay and ageing hallmarks. (**A,B**) Representative images of T1 axons in the medulla of 28-30 old flies labelled with GFP-tagged α-tubulin (tubulin, greyscale and magenta in insets) and the plasma membrane marker myr-Tom (myr, green in insets). Aged neurons in the absence (A) or presence of combined Tau and Shot knockdowns (*Tau_RNAi_ Shot_RNAi_* in B) are compared. Combined Tau and Shot knockdowns enhance phenotypes in ageing neurons, comprising axon swellings often displaying MT unbundling (arrow heads) and MT breaks (asterisks); boxed areas shown as 2-fold magnified double/single-channel images below. (**C**) Quantifications of phenotypes shown in A and B plus conditions of further individual knockdowns for Tau (*Tau_RNAi_*), Shot (*Shot_RNAi_*), and EB1 in an EB1 heterozygote background (*EB1_RNAi_; EB1*^+/Df^); specific knockdowns are indicated below X-axes. Data points are shown as blue circles and as mean ± SEM; P-values obtained via Mann-Whitney tests are indicated above. Data were taken from a minimum of 14 specimens per age group and condition. For detailed statistical values see table S9. Scale bar in A represents 10 μm in all images.

Tau and Shot were shown previously to redundantly regulate MT stability in developing neurons (Voelzmann et al., 2016b). In addition, Shot provides guidance of MT growth (Alves-Silva et al., 2012). We therefore carried out T1-specific knock-down of *shot* either alone or in double knock-down with *tau*, and found a similar increase in MT phenotypes and axonal swellings as observed upon knock-down of *tau* alone, with MT breaks and axonal swellings being clearly enhanced upon double knock-down (Fig. 6C). We extended our analysis to synaptic terminals, revealing a clear exacerbation of deficits in the number of synaptic MTs and an increase in swollen and broken synapses in T1-specific double knock-down (Fig. S6A, C and E).

In contrast to *tau* and *shot*, T1-specific knock-down of *EB1* led to no detectable MT phenotypes (not shown), which contrasts with previous findings in cell culture (Alves-Silva et al., 2012, Hahn et al., 2021). A possible explanation for the lack of *EB1*-deficient phenotypes could be the long half-life of EB1 protein (Alves-Silva et al., 2012) in combination with a potentially incomplete knock-down. To overcome this, we performed T1-specific knock-down of *EB1* in an *EB1* heterozygous mutant background (using an allele carrying a deficiency of the *EB1* gene *Df(2R)Exel6065*). This condition led to an increase in MT breaks, areas of MT disorganisation and the development of axonal swellings in aged axons (Figs. 6C).

None of the knock-down experiments induced MT breaks and unbundling/disorganisation or axonal swellings in young specimens (Fig. S7) suggesting MT phenotypes and axonal swellings are promoted by age in adult specimens.

Taken together, we show that the loss of MT regulator activities such as Tau, Shot and EB1 exacerbate MT phenotypes in aged neurons, suggesting these factors are contributing to the maintenance of axonal MT bundles. Furthermore, their deficient function greatly exacerbates age-associated axonal and synaptic atrophy, suggesting that they are likely factors implicated in the normal process of neuronal ageing.

### Protecting and enhancing MT bundles ameliorates axonal ageing phenotypes

The correlation of MT bundle decay with a reduction in the MT localisation of key MT regulators (Fig. 5), together with their loss of function phenotypes (Fig. 6 and S6) and the late onset of non-MT-related axonal phenotypes (Fig. 1 and 2), might suggest that aberrant MT regulation is a major cause for axonal decay. We therefore reasoned that we may be able to slow down age-related axonal atrophy by protecting MTs from decay during ageing or artificially boosting the function of specific MT regulators.

Counteracting the loss of Tau from MTs by overexpressing Tau, might be counterproductive as suggested by detrimental outcomes of this strategy in *Drosophila* neurons (Ubhi et al., 2007). We therefore decided to focus on EB1 and increased its levels by overexpressing EB1 in T1 neurons. When EB1-expressing brains were visualised at 4-5 weeks, we observed an improvement of all tested ageing phenotypes (Fig. 7D and S6E), namely, reduced disorganisation and less breaks of axonal MT bundles, and the MT bundle diameters and quantity of MTs at the synaptic terminals were significantly increased. In addition, EB1 overexpression led to the improvement of non-MT-related phenotypes, including a reduction in axonal swellings, an increase in axonal calibres and a rejuvenated morphology of synaptic terminals.

**Figure 7:**
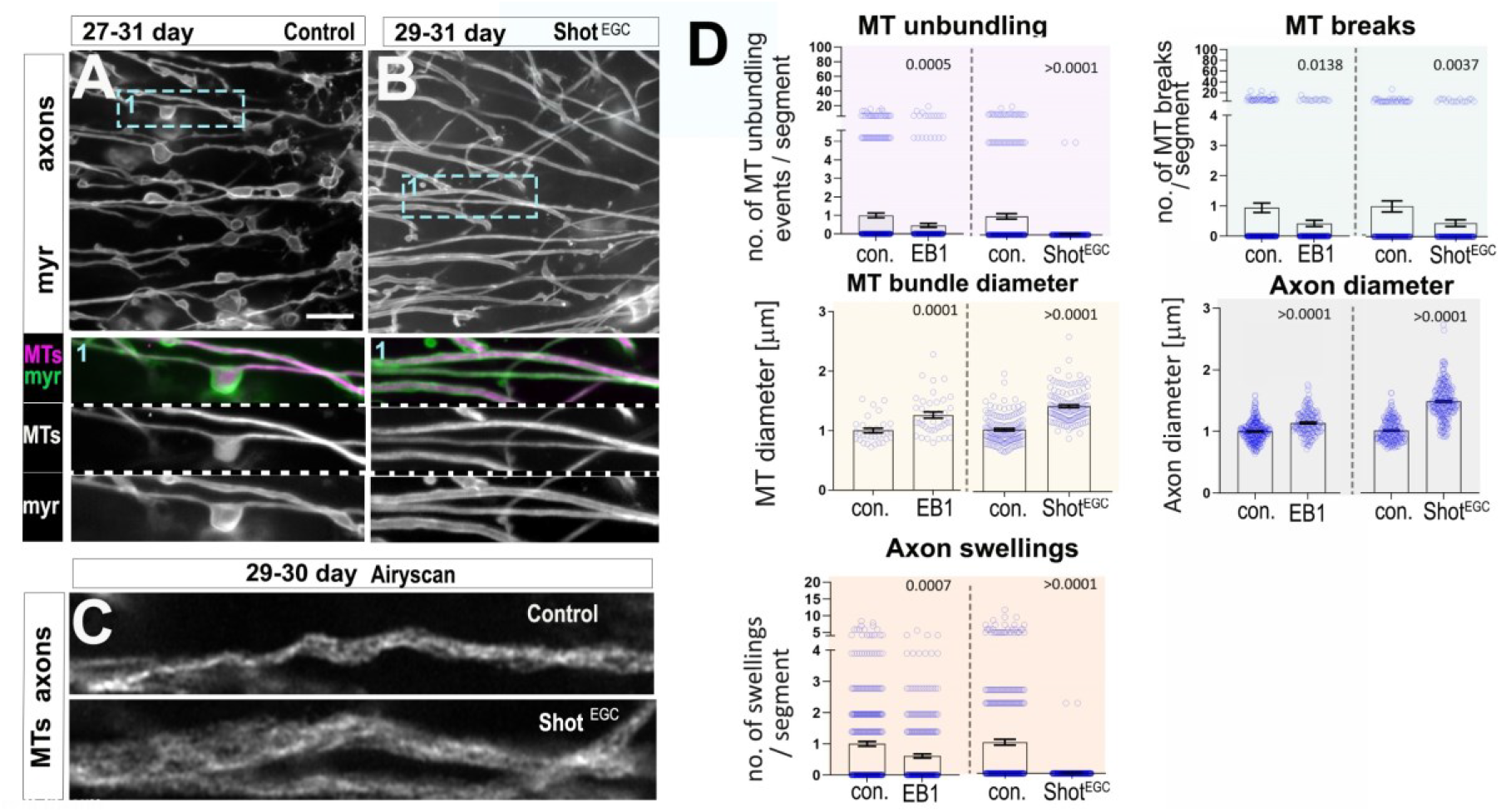
Expression of EB1 and of Shot^EGC^ ameliorates axonal ageing phenotypes. (**A,B**) T1 axons in the medulla of old specimens of 29-31 days old, are labelled with the plasma membrane marker myr-Tom (myr, greyscale in A and B, and green in double-channel images). Ageing phenotypes including axon swellings, axon thinning and MT unbundling can be observed in old specimens in A, but are absent upon T1 specific expression of Shot^EGC^ encoding the C-terminus of Shot (A compared to B, boxed area 1 shown as 2-fold magnified inset where MTs are labelled with GFP-tagged α-tubulin in A or GFP-tagged Shot^EGC^ in B). (**C**) MT axonal bundles from T1 neurons from old specimens (35-38 day old, labelled with GFP-tagged α-tubulin or GFP-tagged Shot^EGC^), imaged using Airyscanning high end confocal microscopy, appear thicker upon Shot^EGC^ expression. (**D**) Quantifications of phenotypes shown in A-C, plus conditions of EB1 ectopic expression. Specific conditions are indicated below X-axes. Data points are shown as blue circles and as mean ± SEM; P-values obtained via Mann-Whitney test are indicated above. Data were taken from a minimum of 13 specimens per group (except for diameter measures of MT bundle with a minimum of 6, and axons with minimum of 10 specimens). For detailed statistical values see table S12. Scale bar in A represents 10 μm in A and B, and 2 μm in C.

Next, we utilised the C-terminus of Shot fused to GFP (Shot^EGC^::GFP) as an alternative to protect MTs from deterioration during ageing. Shot^EGC^ contains both the Gas2-related domain/GRD, which weakly associates with MT lattices and protects them against depolymerisation, and the unstructured, positively charged C-tail which also shows modest association with MT lattices and contains EB1-binding SxIP motives. Combined in the Shot^EGC^::GFP construct, these domains mediate strong association all along MT lattices, fully decorate MT bundles protecting them against depolymerisation and facilitate an association with EB1 (Alves-Silva et al., 2012, Voelzmann et al., 2017). Consistent with our previous work in primary neurons and fibroblasts (Alves-Silva et al., 2012), Shot^EGC^::GFP decorates axonal MTs also in T1 neurons (Fig. 7B). Similar to EB1 overexpression, Shot^EGC^::GFP primarily improved deteriorating MTs and substantially improved all measured ageing phenotypes in axons (Fig. 7A-D) and in synaptic terminals (Fig. S6 B, D and E). Clearly, the strengthening of MT networks via EB1::GFP and Shot^EGC^::GFP expression had a rejuvenating effect on T1 neurons.

When restricting EB1 and Shot^EGC^ overexpression exclusively to mature neurons in the adult brain (using the Gal4/Gal80^ts^ system), all tested age-related MT phenotypes, together with axonal swellings, were significantly improved by Shot^EGC^, whereas EB1 showed a clear positive trend, but mostly not significant compared to control old specimens (Fig. S8). This might suggest that the protective effects mediated by EB1 during ageing requires expression at an early stage, whilst Shot^EGC^ expression in adults is sufficient to block MT and axonal decay.

These findings further support our hypothesis that MT decay can be an important cause for axon and synaptic decay. They further highlight the prospect of employing MT regulation as a target for potential therapeutic intervention to combat the decay of neurons during ageing.

### Shot^EGC^::GFP expression improves the motor performance of flies

To assess whether the improvement of subcellular neuronal features has an impact on the systemic performance of flies, we assessed fly locomotion as a readout known to be affected by ageing (Rhodenizer et al., 2008, White et al., 2010). For this we used negative geotaxis, which is the natural response of flies to walk upwards after being tapped to the bottom of a container; it is quantified by measuring the distance flies climb as a function of time. For our experiments, we used the pan-neuronal *elav-Gal4* driver in combination with the Gal4/Gal80^ts^ system to express EB1::GFP and Shot^EGC^::GFP in all neurons, but restricted it to the adult stage (starting after eclosure). We found that young flies at 4-5 DAE expressing Shot^EGC^::GFP performed similar to age-matched wild-type controls, whereas EB1::GFP expression seemed to be causing a slight decrease in locomotion (Fig. 8). Compared to the young flies, older flies (25-26 DAE) showed a stark decrease in locomotion, and this was partially rescued by the expression of Shot^EGC^::GFP but not of EB1::GFP (Fig. 8).

**Figure 8:**
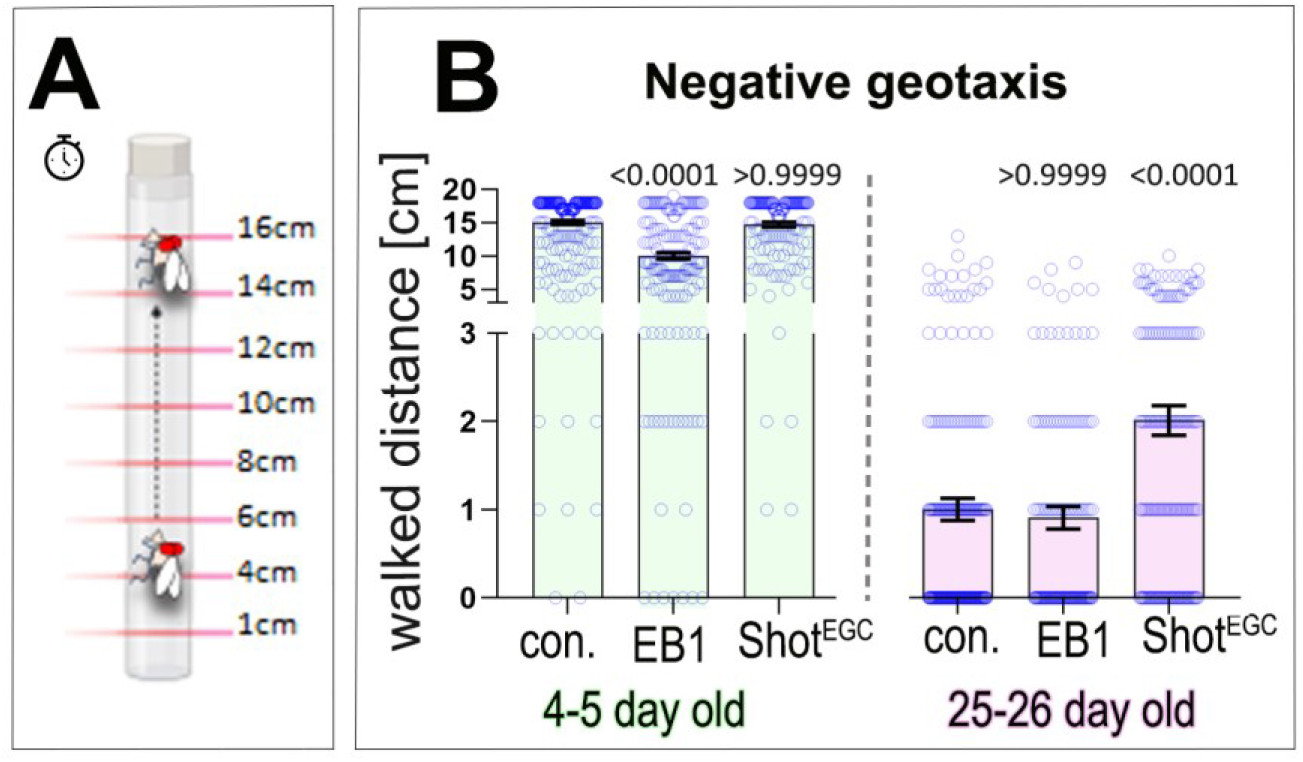
The decline in locomotion of aged flies improves upon adult expression of Shot^EGC^. (**A**) Representation of the negative geotaxis walking assays with flies allowed to walk upwards for 15 seconds, in a narrow graduated cylinder. (**B**) Quantifications of the distance walked by flies in 15 seconds at two different ages 4-5 days old and 25-26 days old. Specific conditions are indicated below X-axes, and includes control flies or flies expressing either Shot^EGC^ or EB1 in adulthood with the pan-neuronal *elav-Gal4* driver using the UAS/Gal4/Gal80^ts^ system (gene expression is induced after development is completed by shifting newly hatched flies from 18 to 29°C, controls are treated with the same regime but are lacking the transgene). Data points are shown as blue circles and as mean ± SEM; P-values obtained via Kruskall-Wallis ANOVA multiple comparison tests are indicated in the graphs; Data were taken from a minimum of 140 specimens per group. For detailed statistical values see table S14.

Given that locomotion requires physiology outside the nervous system likewise affected by ageing, such as the activity of muscles, a full rescue cannot be expected. Nevertheless, our data strongly suggest that restoring neuronal MT networks improves nervous system health with a positive impact on wider body physiology. Therefore, carefully designed strategies targeting MT regulation might provide a path to ameliorate age-related pathological changes in neurons and improve their functionality.

## Discussion

In this study we established a novel model to study neuronal ageing in the *Drosophila* brain. This model is highly accessible to genetic and experimental manipulations, and it displays well-conserved hallmarks of ageing in only 5 weeks. These age-related morphological changes in *Drosophila* axons and synapses, reminiscent of alterations described in higher organisms, include the appearance of axonal swellings, decrease in axonal calibre and the breakdown of synaptic terminals accompanied by membrane swellings or oedemas (Fiala et al., 2007, Ceballos-Baumann et al., 1999, Verdú et al., 2000, Borzuola et al., 2020, Samuel et al., 2011, Rybka et al., 2019, Burke and Barnes, 2006, Fan et al., 2018). Our studies further show that MT deterioration due to decreased turnover is an important change that occurs before the other signs of ageing pathology, that it is likely caused by an age-dependent reduction in the function of MT regulators. Importantly, we showed that reducing key MT regulators such as Tau, EB1 and Shot exacerbates ageing-associated MT defects as well as axonal and synaptic atrophy. As an additional important outcome, our work proved that protecting and enhancing MTs in mature neurons prevents age-related axonal and synaptic deterioration. This MT-based approach also improved neuronal functionality, which naturally declines during ageing.

We propose therefore that MT aberrations can be considered an important causal factor of age-induced neuronal decay, providing new mechanistic insight into neuronal atrophy during normal ageing. Our work also indicates the possibility of employing MT-focussed strategies to protect neurons from deteriorating during ageing.

### Neuronal ageing hallmarks and their conservation with higher organisms

Our studies revealed that the number of T1 neurons is not altered during ageing (Fig. S2). Instead, their axons and synaptic terminals experience remarkable morphological alterations over time (Fig. 1). Overall, axons became thinner and developed swellings, and synaptic terminals appeared to swell and fractionate. Similar morphological alterations have been described in several brain regions from aged rodents and primates. For instance, synaptic terminals in the rat hippocampus and mouse cerebellum develop swellings or oedemas with age (Rybka et al., 2019, Burke and Barnes, 2006, Fan et al., 2018). Furthermore, axonal swellings, or diverticula, were found in axons within the dorsolateral prefrontal cortex of aged rhesus monkey brains (Fiala et al., 2007), and the diameter of axons within the mouse peripheral nervous system was found to decrease with age (Ceballos-Baumann et al., 1999, Verdú et al., 2000, Hamilton et al., 2016, Borzuola et al., 2020). Critically, such biological changes that develop naturally during ageing in higher organisms, appeared to occur in the absence of neuronal loss (Franciscovich et al., 2008) and correlated with functional decay of neuronal networks (Brickman et al., 2007, Koini et al., 2018). Indeed, axonal swellings are considered a hallmark of axonal degeneration and axonal atrophy (Wang et al., 2020), and axonal calibre/diameter is a key determinant of the conduction properties of neurons, with thinner axons correlating with decreased conductivity (Fan et al., 2017). Therefore, the presented herein age-dependent alterations in *Drosophila* neurons are consistent with those observed in mammals, proving the validity and efficacy of our model.

### The importance of the MT cytoskeleton in neuronal ageing

Although none of the previous studies linked the aforementioned morphological alterations to defects in the MT cytoskeleton, MTs have been proposed to be affected by ageing in other invertebrates models (Pan et al., 2011) and in higher organisms (Kounakis and Tavernarakis, 2019). Reduction in MT density and alterations in MT organisation within axons are both reported to take place within human and non-human primate brains during ageing (Cash et al., 2003, Fiala et al., 2007). In human pyramidal neurons, a 55% reduction in MT density was reported prior to the formation of Tau neurofibrillary tangles (Cash et al., 2003), suggesting that changes occur at the level of the cytoskeleton before other bona fide ageing hallmarks. In *C. elegans*, MT bundles are disorganised in aged specimens (Pan et al., 2011). Considering the previous reports together with our new findings, we propose that these changes in the MT cytoskeleton observed in aged neurons constitute a widespread phenomenon, apparent in primates and invertebrates.

The work presented here further demonstrates that there is a causative link between MT deterioration during ageing and natural axonal decay exemplified by the formation of axonal swellings and the decrease in axonal calibre. We show not only that MT bundle impairment precedes other neuronal morphological changes, but also that decreasing or enhancing the function of specific MT regulators is sufficient to aggravate or rescue axonal decay respectively. However, the mechanisms by which MT deterioration leads to perturbations of the axon are yet not understood, but may involve changes in membrane tension and/or the axonal membrane associated periodic skeleton, both of which are highly dependent on the MT cytoskeleton (Zhong et al., 2014, Qu et al., 2017, Datar et al., 2019). For example, compromising MT integrity generates membrane tension-driven instabilities, leading to axonal beading in cultured neurons (Jaworski et al., 2009, Kopf et al., 2020, Datar et al., 2019). MT breaks or areas of disorganisation could also halt transport of organelles along the axon, leading to acute accumulations and the development of swellings. In fact, disruptions in axonal transport and aberrant accumulation of organelles along the axon are frequently observed in aged organisms from different species, including rhesus monkey, *Drosophila* and *C. elegans* (Fiala et al., 2007, Vagnoni and Bullock, 2018, Vagnoni et al., 2016, Li et al., 2016). In *C. elegans*, ageing affects the activity of the motor protein UNC-104, preventing its localisation in axons, and rendering neural circuits more vulnerable (Li et al., 2016). We previously found that the loss of Tau and Shot similarly affects the translocation of UNC-104 to axons. Over time, this leads to a decrease of key synaptic proteins in synapses (Voelzmann et al., 2016b). The fact that MT deterioration causes synaptic decay and axonal swellings in our model may also suggest that transport capacity in T1 neurons also declines with age.

Importantly, we found that by preserving MTs during ageing, by the means of increasing the function of specific MT regulators, we could improve axon’s health (Fig. 7), thus firmly demonstrating that MTs are essential drivers of neuronal decay during ageing. Indeed, in neurodegenerative diseases (Alzheimer’s disease, fronto-temporal dementia, Parkinson’s disease, amyotrophic lateral sclerosis, hereditary sensory autonomic neuropathy), significant decay of axonal MT bundles is observed (Adalbert and Coleman, 2013, Cash et al., 2003, Edvardson et al., 2012, Eyer et al., 1998, Seehusen et al., 2016). It remains an open question how these diseases cause such alterations, and whether MT decay is occurring in parallel to axonal and synaptic atrophy. However, our results suggest that MT deterioration could be a mediator. Ageing is considered a key risk factor common to the aforementioned neurodegenerative diseases, and common to both ageing and these pathologies are MT deterioration, axonal swellings, synaptic fragmentation and defective transport (Sleigh et al., 2019, Fiala et al., 2007, Vagnoni and Bullock, 2018a, Vagnoni et al., 2016, Salvadores et al., 2017, Cash et al., 2003, Brunden et al., 2017, Kounakis and Tavernarakis, 2019, Sferra et al., 2020). It may be that MT bundle deterioration is a key shared mechanism that when unrectified, can trigger transport deficits leading to axonal and synaptic decay. In this model, age would render the maintenance of MT bundles less efficient, ultimately sensitising axons, making them even more vulnerable in disease conditions.

### MT regulation in mature neurons

How axonal MT bundles are maintained in mature neurons and during ageing has gained limited attention in the past. Our tubulin::GFP pulse-chase experiments suggest that there is continued MT turnover in mature axons which may involve MT disassembly, nucleation, polymerisation and repair (Kleele et al., 2014, Triclin et al., 2021), yet MT turnover appears severely diminished with age. The observed MT breaks in axons and a decrease in the diameter of MT bundles with age might therefore be the direct consequence of net loss of MT volume, due to an increase in MT instability causing MT loss, or a decrease in the rate at which MTs are replaced.

A hint pointing to the mechanisms mediating loss of MT volume during ageing is suggested by our findings that attenuating the function of Tau, EB1 and the spectraplakin Shot, exacerbates ageing-associated MT breaks in axons from adult *Drosophila*. These findings suggest that the regulation of MT turnover in axons from adults involve functions of Tau, EB1 and Shot. Tau and Shot ability to promote MT stability (Voelzmann et al., 2016) as well as EB1 roles in promoting MT polymerisation dynamics and the repair of damaged MT shafts (Triclin et al., 2021, Alves-Silva et al., 2012, Elliott et al., 2005, Li et al., 2011, van Haren et al., 2018) could directly impact the rate at which MTs are lost and replaced in axons of mature neurons during ageing.

Along with the loss of MT volume in aged specimens, we also observed foci of disorganised MTs in mature axons, which are increased in number by reducing the function of Tau, EB1 and Shot. Such areas could arise from a decrease in MT bundling caused by loss of Tau during ageing, since Tau can bundle MTs by the means of its projection domain (Chung et al., 2016). Similarly, Shot and other spectraplakin are proposed to bundle MTs through mechanisms not yet understood (Alves-Silva et al., 2012, Voelzmann et al., 2017). Alternatively, the loss of linkage of polymerising MTs to cortical actin cytoskeleton can cause aberrant MTs extension into disorganised and curved conformations contributing to disorganised MT foci in aged axons. Shot, EB1 and Tau cooperate in this linkage function in developing neurons (Alves-Silva et al., 2012, Hahn et al., 2021, Sanchez-Soriano et al., 2009, Voelzmann et al., 2017, Voelzmann et al., 2016). This function appears to remain essential during ageing as suggested by our data showing an increase in disorganised MT foci during ageing upon Tau, Eb1 and Shot loss, and the rescue of these phenotypes after expression of EB1 and Shot^EGC^.

Our studies suggest that MT deterioration is likely due to a natural age-dependent reduction in the function of MT regulators such as Tau and EB1. It remains to be established what causes such reduction, but altering their posttranslational modification state is a likely explanation (Ramkumar et al., 2018, Nehlig et al., 2017), or they may be impacted by changes in cytoplasmic ROS or Calcium (Goldblum et al., 2020, Kapur et al., 2012), all known to change in the ageing brain. Our findings may once more have important parallels to the aged human brain where the function and localisation of Tau is also altered (Saha and Sen, 2019, Schöll et al., 2016).

### The value of the MT cytoskeleton as therapeutic target

In this work we showed that improving MT’s health halts neuronal decay during ageing. By targeting and protecting MTs, we may be able to decrease susceptibility to age-related diseases for instance by decreasing the age-related decline in axonal transport capacity. The use of such MT-targeting strategies to battle ageing, may also be of benefit in other contexts, as comparable decline in MTs might be occurring in other cell types (and/or tissues). For instance, altered MT dynamics during spindle formation is a primary cause for age-related chromosome segregation errors in oocytes. Additionally, changes in mitochondria activity observed in aged mouse oocytes are linked to alterations in the cytoskeleton, and together, could contribute to increased infertility in oocytes from female mammals with age (Nakagawa and FitzHarris, 2017, Kim et al., 2022), thus highlighting the potential of targeting MTs to slow down the ageing process in different cell types.

Several studies including ours have pointed to the validity of this approach. For example, targeting MTs by increasing their acetylation, which is frequently linked to MT stability, protects axons and transport of mitochondria in a mouse trauma model caused by intracerebral haemorrhage (Yang et al., 2022). Furthermore, manipulation of MTs by the use of MT-targeting agents (MTAs) has proven valuable in the context of age-related neurodegenerative diseases, as suggested by recent reports highlighting the beneficial therapeutic effects of MT-stabilising compounds such as Epothilone D in several models of tauopathies (Varidaki et al., 2018, Fernandez-Valenzuela et al., 2020, Wordeman and Vicente, 2021). However, MTAs can induce toxicity and trigger sides effects such as neuropathy, and their prolonged usage can lead to drug resistance (Wordeman and Vicente, 2021). Therefore, modifying MTs by manipulating MT-binding proteins may be a promising alternative approach (Llorente-González et al., 2021, Ramkumar et al., 2018).

## Material and Methods

### Fly stocks and husbandry

The following fly stocks were used in this project. Gal4 driver lines include: *elav-Gal4* (3^rd^ chromosomal, expressing pan-neuronally at all stages; (Luo et al., 1994)), *GMR31F10-Gal4* (Bloomington #49685; expressing in T1 medulla neurons; (Qu et al., 2019), *GMR53G02-GAL4* (Bloomington #50446; expressing in T2 medulla neurons; (Tuthill et al., 2013)). Mutant stocks: EB1 deficiency *Df(2R)Exel6065* (EB1^*Df*^; Bloomington stock #7547). Lines for transgene targeted expression were *UAS-Shot^EGC^-GFP* and *UAS-EB1-GFP* (Alves-Silva et al., 2012), *UAS-GFP-α-tubulin84B* (Grieder et al., 2000), *UAS-RedStinger* (Bloomington stock #84277) and *UAS-myr::tdTomato* (Bloomington stock #32222). Lines for targeted gene knockdowns were *UAS-tau^RNAi^ (tau^GD25023^* Vienna *Drosophila* RNAi Center, Austria (Bolkan and Kretzschmar, 2014)), *UAS-shot^RNAi^* (Subramanian et al., 2003) and *UAS-EB1^RNAi^ (EB1^24451^* Vienna *Drosophila* RNAi Center, Austria, (Alves-Silva et al., 2012).

For ageing experiments, all flies were maintained during ageing at low density (maximum of 20 flies per vial) on standard sugar-yeast molasses medium, at 29°C unless otherwise specified. Experimental flies were transferred to fresh vials every 3 days.

### Dissection of adult fly heads

Dissections of *Drosophila* brain were performed in Dulbecco’s PBS (Sigma, RNBF2227) after briefly anaesthetising the specimens on ice. For live imaging, dissected brains with their laminas and eyes attached were placed into a drop of Dulbecco’s PBS on MatTek glass bottom dishes (P35G1.5-14C), with a spacer and covered by coverslips. Brains were immediately imaged.

### Immunohistochemistry

*Drosophila* brain dissections were performed as explained above and fixed in 4% paraformaldehyde (PFA; in 0.05 M phosphate buffer, pH 7–7.2) for 30 min at room temperature (RT). For anti-EB1 staining, brains were fixed at −20°C for 10 mins in +TIP fix (90% methanol, 3% formaldehyde, 5 mM sodium carbonate, pH 9; stored at −80°C (Hahn et al., 2021a)). Brains were washed in PBT (PBS with 0.3% Triton X-100). Antibody staining and washes were performed with PBT.

Antibodies used include: anti-DmEB1 (gift from H. Ohkura; rabbit, 1:2000; (Elliott et al., 2005)); anti-GFP (ab290, Abcam, 1:500); FITC-, Cy3- or Cy5-conjugated secondary antibodies (Jackson ImmunoResearch); and STAR 580 (Abberior STAR 580) for super resolution. Specimens were embedded in Vectashield (VectorLabs).

### Microscopy and data analysis

For all ageing parameter, *Drosophila* brains were imaged at the Centre for Cell Imaging at the University of Liverpool with a 3i Marianas Spinning Disk Confocal Microscope, except for the study of MT diameters where the STED system STEDYCON or the LSM 900 Airyscan2 was used.

To measure ageing hallmarks in the optic lobe of adult flies (except for MT diameter), brains were dissected as explained above and immediately imaged with a 3i Marianas Spinning Disk Confocal Microscope at the Centre for Cell Imaging at the University of Liverpool. A section of the medulla columns comprising the 4 most proximal axonal terminals was used to quantify MTs axonal phenotypes and the number of swellings. To measure Tub-GFP incorporation into axonal MTs, a line was drawn along these axons, using the segmented line tool in Fiji, in order to measure the mean intensity. To measure the intensity from UAS-myr::tdTomato, a fitting box around the axon was drawn using the polygon selection tool and the mean intensity calculated. To measure axonal and MT core diameter, brain images were orientated so the columns of medulla axonal projections were displayed horizontally. The line tool of ImageJ was used to measure the diameter of axons and their MT core, at evenly spaced unbiased positions predetermined by a grid. A mean per axon from multiple measure points was obtained. To score synaptic readouts, a square was drawn through the central part of the medulla and synapses within scored (between 30-50 per medulla). Synaptic MTs and morphological categorization were manually calculated. Nuclei labelled with RedStinger were quantified manually in Fiji, Image J software.

For pixel intensity analysis of Tau in axons, maximal projections were generated from Z-stacks using Fiji, Image J software. With the line tool, 4 regions of interest (ROI) per medulla, spanning across 8 axon bundles, were positioned parallel to each other. Intensity histograms were generated per ROI and pixel intensity threshold bins for grey values >350 were set. Peaks above the threshold were counted and the mean was taken across the 4 ROIs. For pixel intensity analysis of EB1 in axons, maximal projections from 6 stacks were generated from two areas of each medulla (Upper & lower half). Ten medulla axon columns were selected using the ‘manual draw’ tool on ImageJ, and mean intensity was measured in the EB1 channel. The mean intensity per medulla was calculated and plotted.

### MT-binding assay and western blots

MT-binding assays were performed as previously discussed (Quraishe et al., 2013, Cowan et al., 2010). In brief, for each age group, 6 heads were homogenised in 40 μl of MT-binding assay buffer (100 mM MES pH 6.8, 500 μM MgSO4, 1 mM EGTA, 4 mM DTT, 2 mM dGTP, 20 μM taxol, 0.1% triton X-100, 30 mM NaF, 20 mM sodium pyrophosphate, 40 mM 2-glycerophosphate, 3.5 mM sodium orthovanadate, 10 μM staurosporine and protease inhibitor cocktail). Homogenates were centrifuged at 12000g for 1 hour at 4°C. The supernatant, representing the cytosolic fraction, was transferred to a fresh tube while the pellet, representing the fraction enriched for MT-bound proteins, was resuspended in 20 μl of the MT-binding assay buffer. Samples were heated in Laemmli buffer for 10 minutes at 95°C and separated by electrophoresis.

### Western blots

For total protein level analysis, heads were homogenised in ice-cold RIPA buffer, supplemented with Protease and Phosphatase inhibitors. The samples were heated in Laemmli buffer for 10 min at 95°C before electrophoresis. For electrophoresis separation, extracts were resolved in 10% Stain-free SDS-PAGE gels (BioRad) in running buffer (25 mM Tris Base, 192 mM Glycine, 10% w/v SDS) and transferred to a nitrocellulose membrane followed by blocking in 5% bovine serum albumin for 1 hour at room temperature. Proteins were probed with primary antibodies and detected by incubation with HRP-conjugated secondary antibodies (Invitrogen). The following primary antibodies were used: anti-GAPDH (Thermofisher GA1R), anti-GFP (Abcam 6673), anti-DTau (L. Partridge) and anti-EB1 (H. Ohkura).

### Negative geotaxis assay

One day prior to the experiment, a maximum of 20 flies were transferred to fresh media vials. On the day of the experiment, flies were transferred to graduated vials without the use of CO_2_. Flies were given 30 minutes to acclimatise to the new environment. After this time, flies were tapped to the bottom of vials and their walking behaviour was filmed. The position of each fly after 15 seconds time was annotated. Three technical replications were conducted which included all together between 220 to 340 young flies and between 170 to 290 old flies per genotype.

### Statistical analysis

Statistical analyses were performed in GraphPad Prism 9 using Mann-Whitney Rank Sum Tests (indicated as P_MW_), or via Kruskall-Wallis ANOVA multiple comparison tests or Chi^2^ (P_Chi_), with 95%confidence intervals. The exact p-values are indicated in the graphs and the exact sample size and other statistical values are indicated in the supplementary tables S1 to S14.

## Acknowledgements

This work was made possible through support to N.S.S by the BBSRC (BB/M007456/1, BB/R018960/1) and to R.G, S.S and N.S.S by the Wellcome Trust (WT204002).

We thank the University of Liverpool Imaging Facility for the help. We thank Hiro Ohkura for kindly providing the DmEB1 antibody. We thank Andreas Prokop for constructive comments on this manuscript. Stocks obtained from the Bloomington *Drosophila* Stock Center (NIH P40OD018537) were used in this study.

---

**Figure S1.**
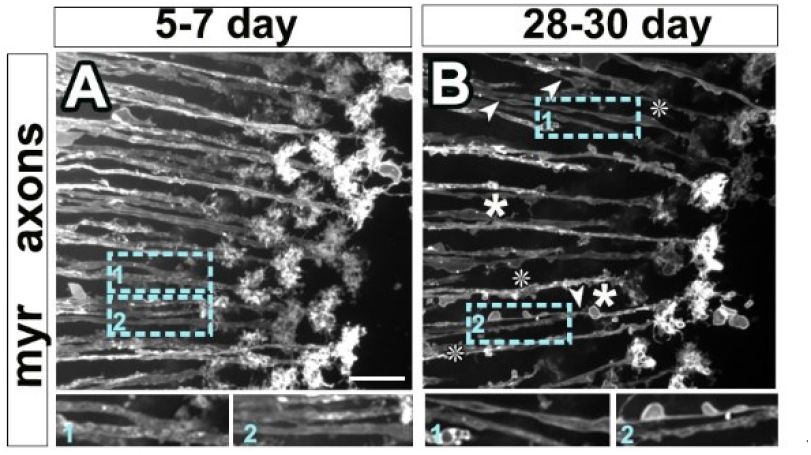
Widespread deterioration of neurons within the *Drosophila* visual system with age. (**A,B**) Axons and terminals of the L2 neurons within the medulla of the optic lobe from young (5-7 day, A) and old specimens (28-30 day, B), labelled with the plasma membrane marker myr-Tom (myr). In aged specimens, axons show thinning (arrow heads) and swellings (asterisks; dashed blue boxes shown as 1.7-fold magnified images below). Scale Bar for A and B can be found in A bottom right (A-B 10 μm).

**Figure S2.**
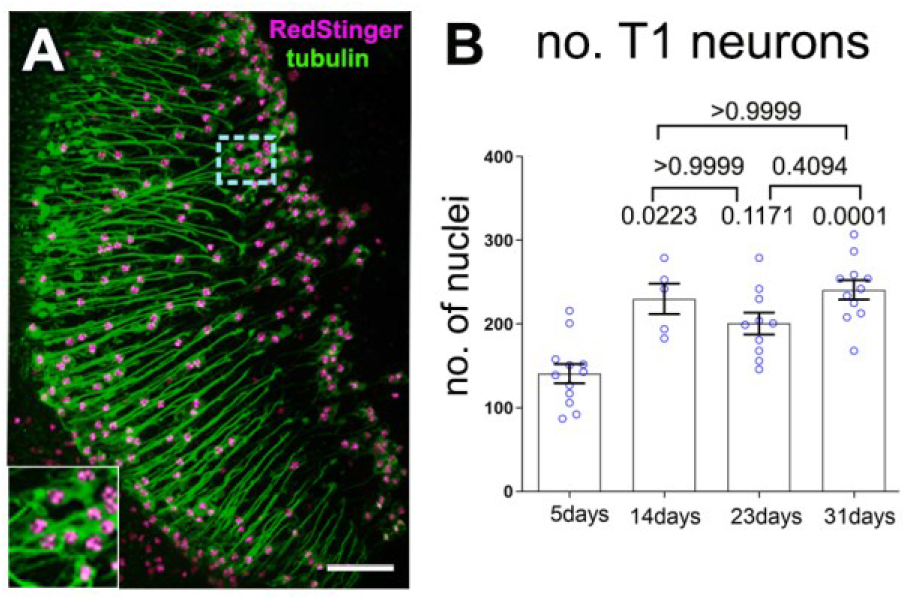
Absence of neuronal death amongst T1 neurons during ageing. (**A**) T1 neurons labelled with the nucleus marker RedStringer (magenta) and GFP-tagged α-tubulin (green). Dashed blue boxes shown as 2-fold magnified image in the inset below). (**B**) Quantification of nuclei labelled with RedStinger at different ages to determine the number of T1 neurons. Data are shown as mean ± SEM of nuclei per medulla with individual data points in blue; p-values obtained with Kruskall-Wallis ANOVA test for the different conditions are indicated in the graph. For detailed statistical values see table S2. Scale Bar for A represents 30μm.

**Figure S3.**
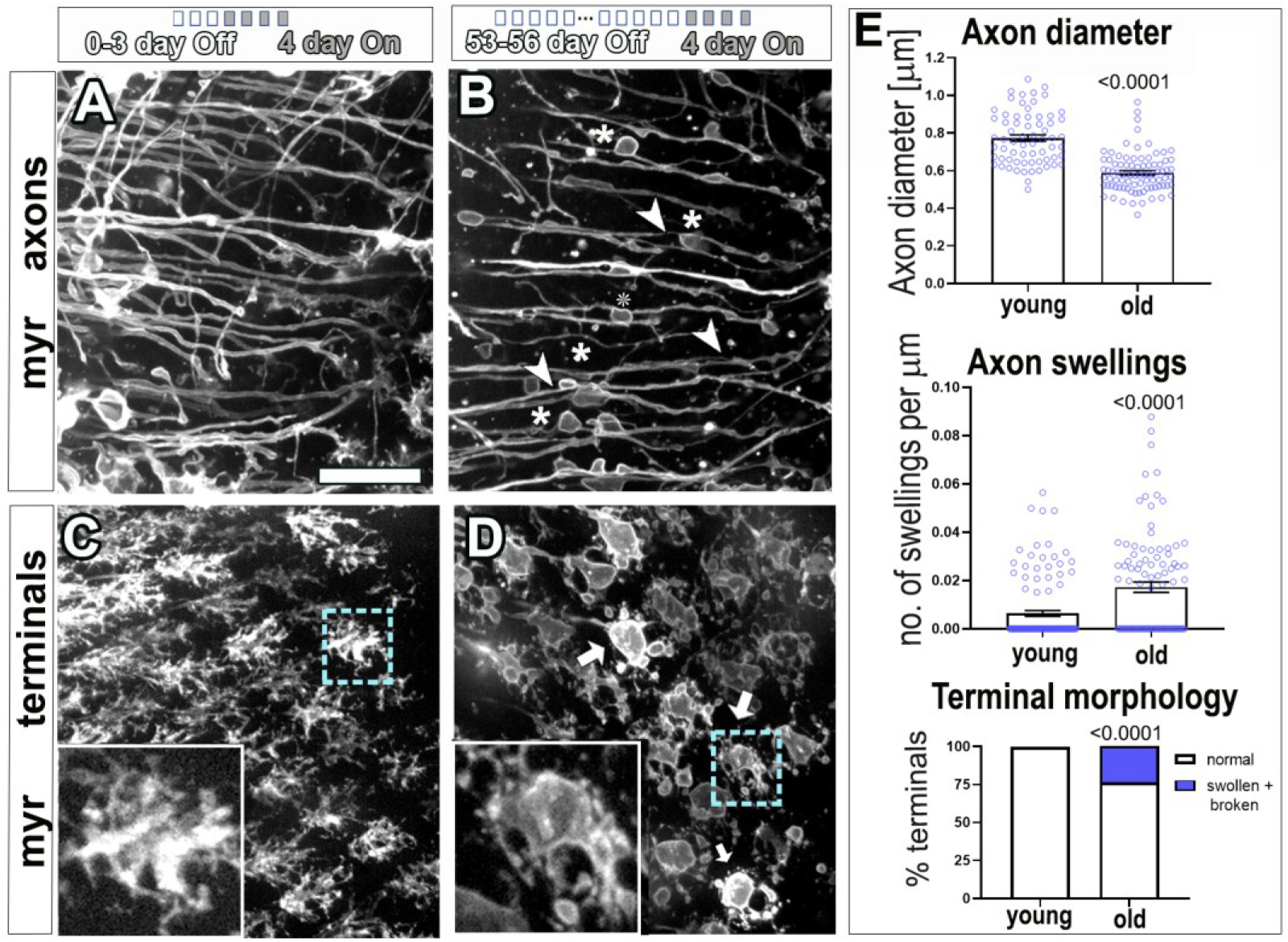
The deterioration of T1 neurons during ageing is independent of temperature and marker expression. **(A-D**) T1 axons (top) and synaptic terminals (bottom) in the medulla of flies labelled with the plasma membrane marker myr-Tom (myr) using the UAS/Gal4/Gal80^ts^ system. Gene expression is induced by the shift of temperature from 18 to 29°C. Flies were kept at 18°C throughout development and adult life until the last 4 days before imaging, at which point they were sifted to 29°C to initiate myr-Tomato gene expression. Young specimens (A,C; 4-7 day old flies with 0-3 days at 18°C “Off” + 4 days at 29°C “On”) are compared to old specimens (B,D; 57-60 day old flies with 53-56 days at 18°C “Off” + 4 at 29°C “On”). In aged specimens, axons show thinning (arrow heads) and swellings (asterisks), and synaptic terminals appear broken down and with swellings (arrows and dashed blue box shown as 3-fold magnified image in the inset below). **E**) Quantifications of phenotypes shown in A-D, with young versus old indicated on the X-axes. In the top two graphs, data points are shown in blue and as mean bars ± SEM (p-values obtained via Mann-Whitney test are indicated above). For terminal morphology, data are shown as distribution of normal versus swollen/broken synapses (significance obtained via Chi-square test indicated above). Data were taken from a minimum of 5 specimens per age group. For detailed statistical values see table S3. Scale bar in A represents 20 μm in all images.

**Figure S4.**
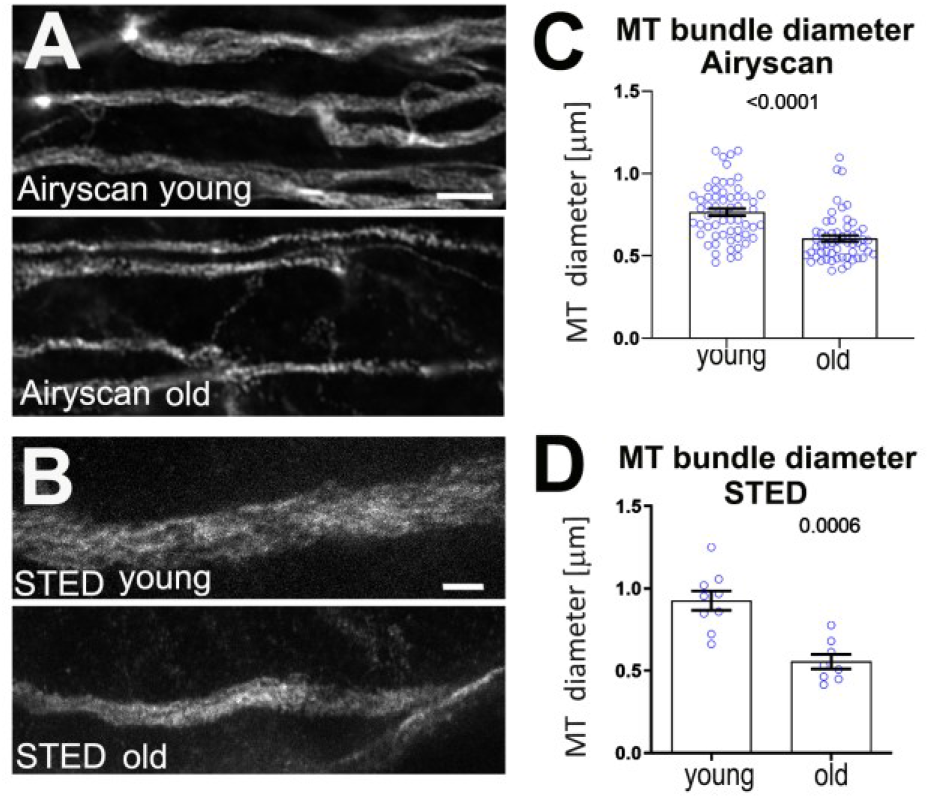
The MT bundle within axons thinners with age. (**A and B**) MT axonal bundles from T1 neurons from young (8 to 10 day old) and old (34-36 day old) specimens labelled with GFP-tagged α-tubulin, are imaged using Airyscanning high end confocal microscopy (A) or stimulated emission depletion (STED) super-resolution microscopy (B). (**C-D**) Images derived from both imaging techniques have been used to quantify the diameter of MT bundles (C with Airyscanning and D with STED). Results are presented as mean ± SEM with individual data points in blue. P-values obtained with Mann-Whitney test are indicated in each graph. Data points were taken from at least 13 specimens per age group for Airy Scanning and 3 specimens for STED. For detailed statistical values see table S4. Scale bar in A represents 4 μm and in B 1 μm.

**Figure S5:**
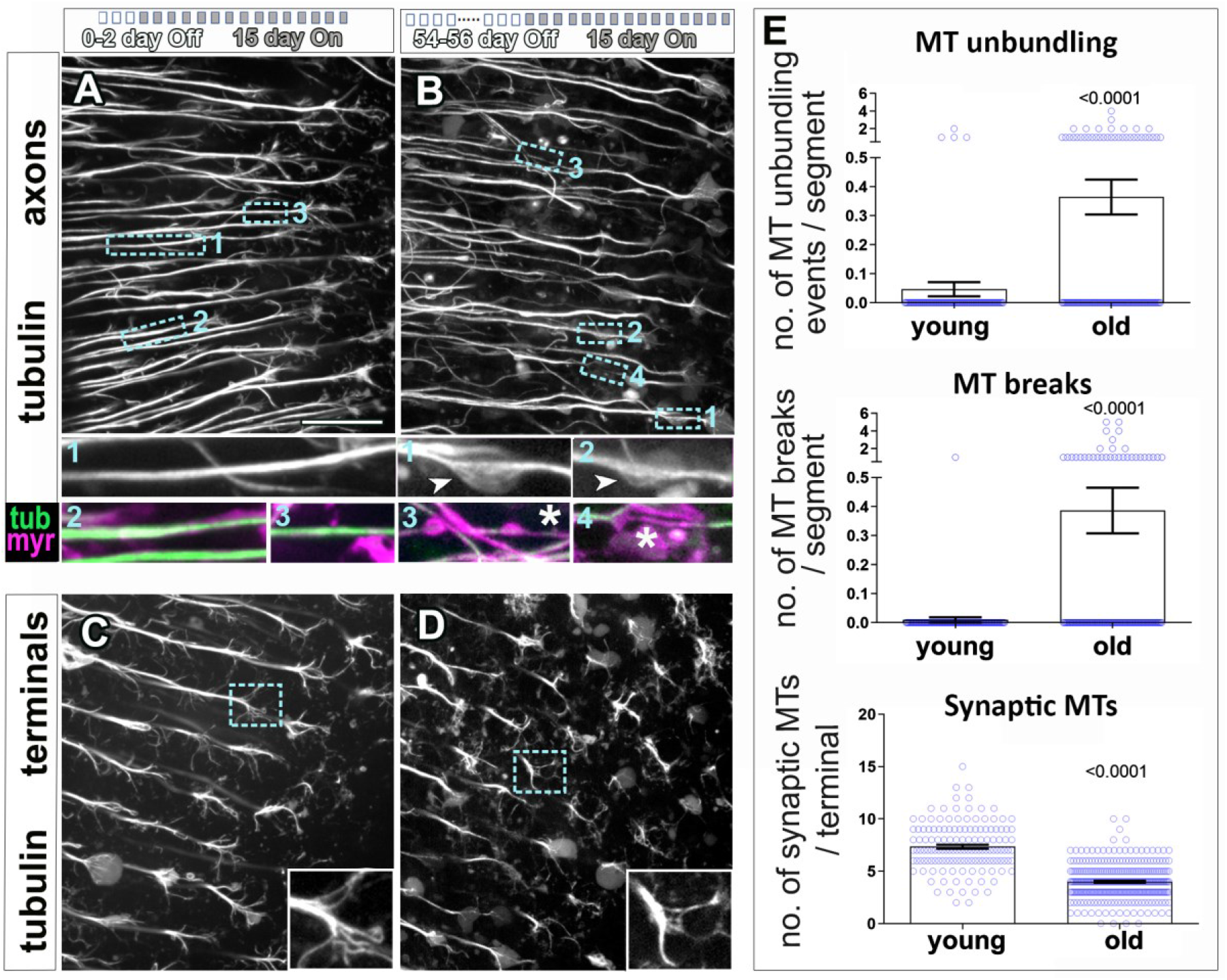
MT alterations during ageing are independent of the MT reporter expression and are specific of aged specimens. (**A-D**) T1 axons (top) and synaptic terminals (bottom) in the medulla of flies with their MTs labelled with GFP-tagged α-tubulin (tubulin, in greyscale images and green in insets) and the plasma membrane marker myr-Tom (myr, magenta), using the UAS/Gal4/Gal80^ts^ system. Gene expression is induced by the shift of temperature from 18 to 29°C. Flies were kept at 18°C throughout development and adult life until the last 15 days before imaging, at which point they were shifted to 29°C to initiate gene expression. Young specimens (A,C; 15-17 day old flies with 0-2 days at 18°C “Off” + 15 days at 29°C “On”) are compared to old specimens (B,D; 69-71 day old flies with 54-56-days at 18°C “Off” + 15 at 29°C “On”). In A and B, cyan encircled boxes are shown 3.7-fold magnified below with old flies in B, showing MT unbundling (insets 1 and 2 in B) as well as breaks and thinning of MTs (insets 3 and 4 in B) compared to young axons. In D, old flies also show a reduction of splayed MTs in synaptic terminals when compared to young controls in C (boxed areas shown as 2-fold magnified insets). (**E**) Quantifications of phenotypes shown in A-D, with young versus old indicated on the X-axes; data points are shown as blue circles and as mean ± SEM; p-values obtained via Mann-Whitney tests are indicated above. Data were taken from a minimum of 5 specimens per age group. For detailed statistical values see table S5. Scale bar in A represents 10 μm, and 25 μm in B.

**Figure S6:**
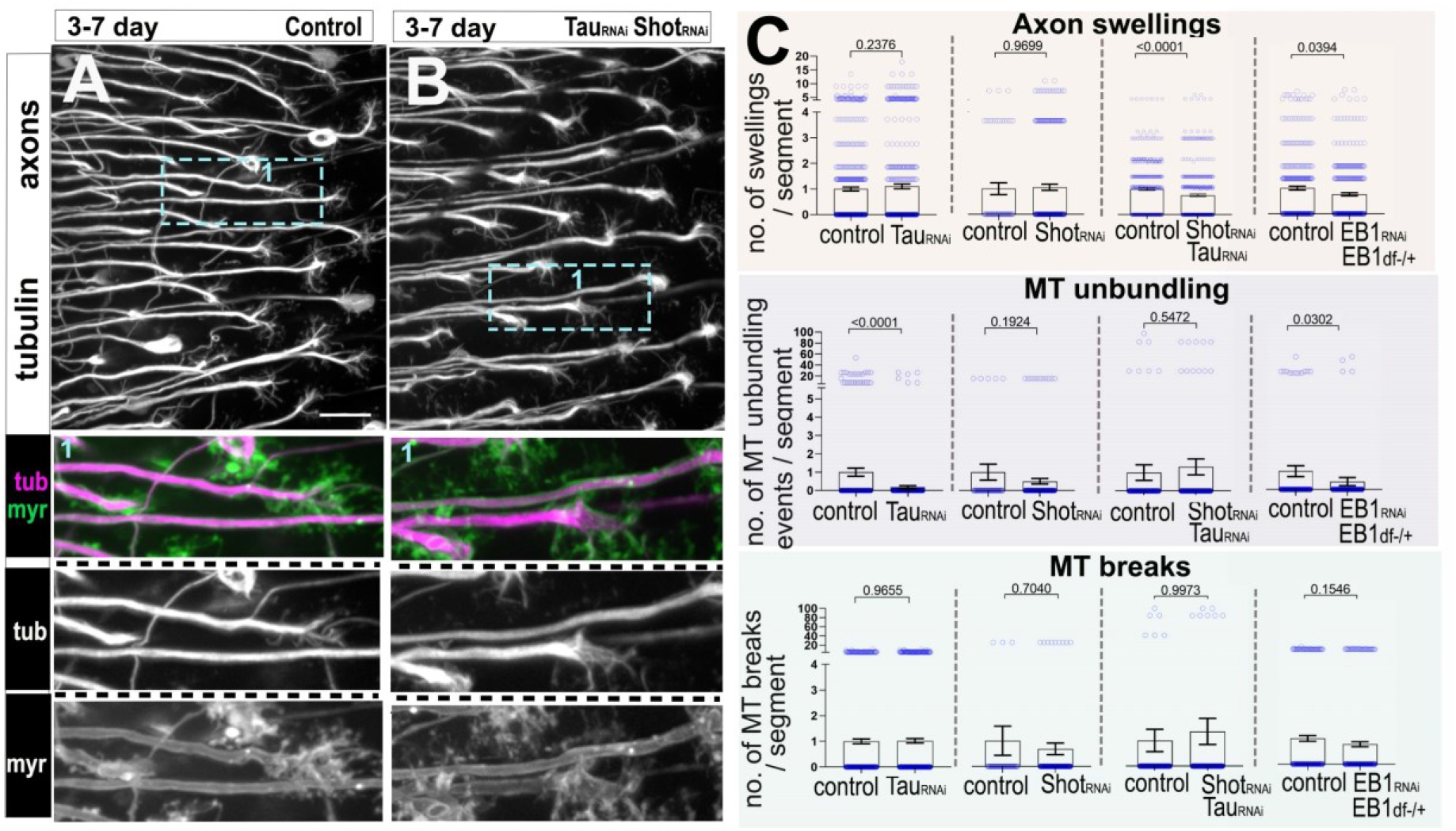
The deterioration of synaptic terminals during ageing can be exacerbated or rescued by altering the function of MT regulators. (**A-D**) Terminals of the T1 medulla projections from old specimens (28-30 day A and C; 29-31 day B and D) with the plasma membrane labelled with myr-Tom (myr) and their MTs with GFP-tagged α-tubulin (A-C) or GFP-tagged Shot^EGC^ (D). Ageing phenotypes at the synaptic terminals including the decrease in synaptic MTs (arrowheads) and the rise in broken and swollen terminals (asterisks) increase in the presence of combined Tau and Shot knockdowns (Tau_RNAi_ Shot_RNAi_ in C) and are suppressed by the expression of Shot^EGC^ (D). (**E**) Quantifications of phenotypes shown in A to D plus conditions of EB1 ectopic expression. Specific conditions are indicated below X-axes. In the upper graphs, data points are shown as blue circles and as mean ± SEM; P-values obtained via Mann-Whitney test are indicated above. For terminal morphology, data are represented as distribution of normal versus swollen/broken synapses; significance obtained via Chi-square test is indicated above. Data were taken from a minimum of 14 specimens per group. For detailed statistical values see table S10. Scale bar in A represents 10 μm in A-D.

**Figure S7:**
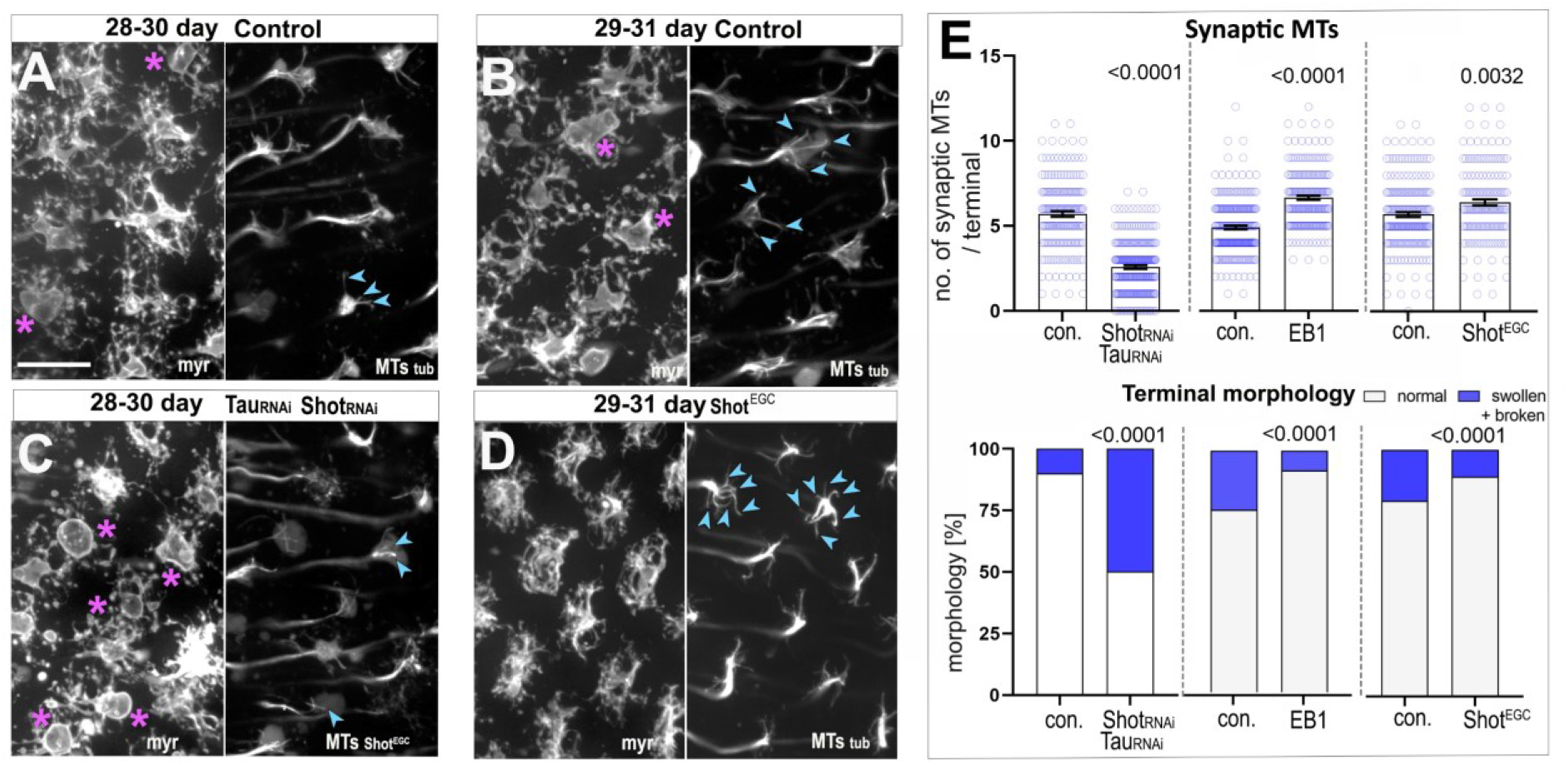
MT decay and ageing hallmarks are not present in young specimens with EB1, Tau and Shot knockdowns. (**A,B**) Representative images of T1 axons in the medulla of 3-7 days old flies labelled with GFP-tagged α-tubulin (tubulin, greyscale and magenta in insets) and the plasma membrane marker myr-Tom (myr, green in insets). Images show specimens in the absence (A) or presence of combined Tau and Shot knockdowns (*Tau_RNAi_ Shot_RNAi_* in B). MTs axonal bundles maintain their diameter and organisation, and axons lack swellings and areas of decreased diameter in the presence of knockdowns; boxed area shown as 2-fold magnified double/single-channel images below. (**C**) Quantifications of phenotypes shown in A and B plus conditions of further knockdowns for Tau (*Tau_RNAi_*), Shot (*Shot_RNAi_*), and EB1 in an EB1 heterozygote background (*EB1_RNAi_ EB1*^+/Df^); specific knockdowns are indicated below X-axes. Data points are shown as blue circles and as mean ± SEM; P-values obtained via Mann-Whitney tests are indicated above. Data were taken from a minimum of 13 specimens per age group and condition with the exception of 8 specimens for Shot. For detailed statistical values see table S11. Scale bar in A represents 10 μm in all images.

**Figure S8:**
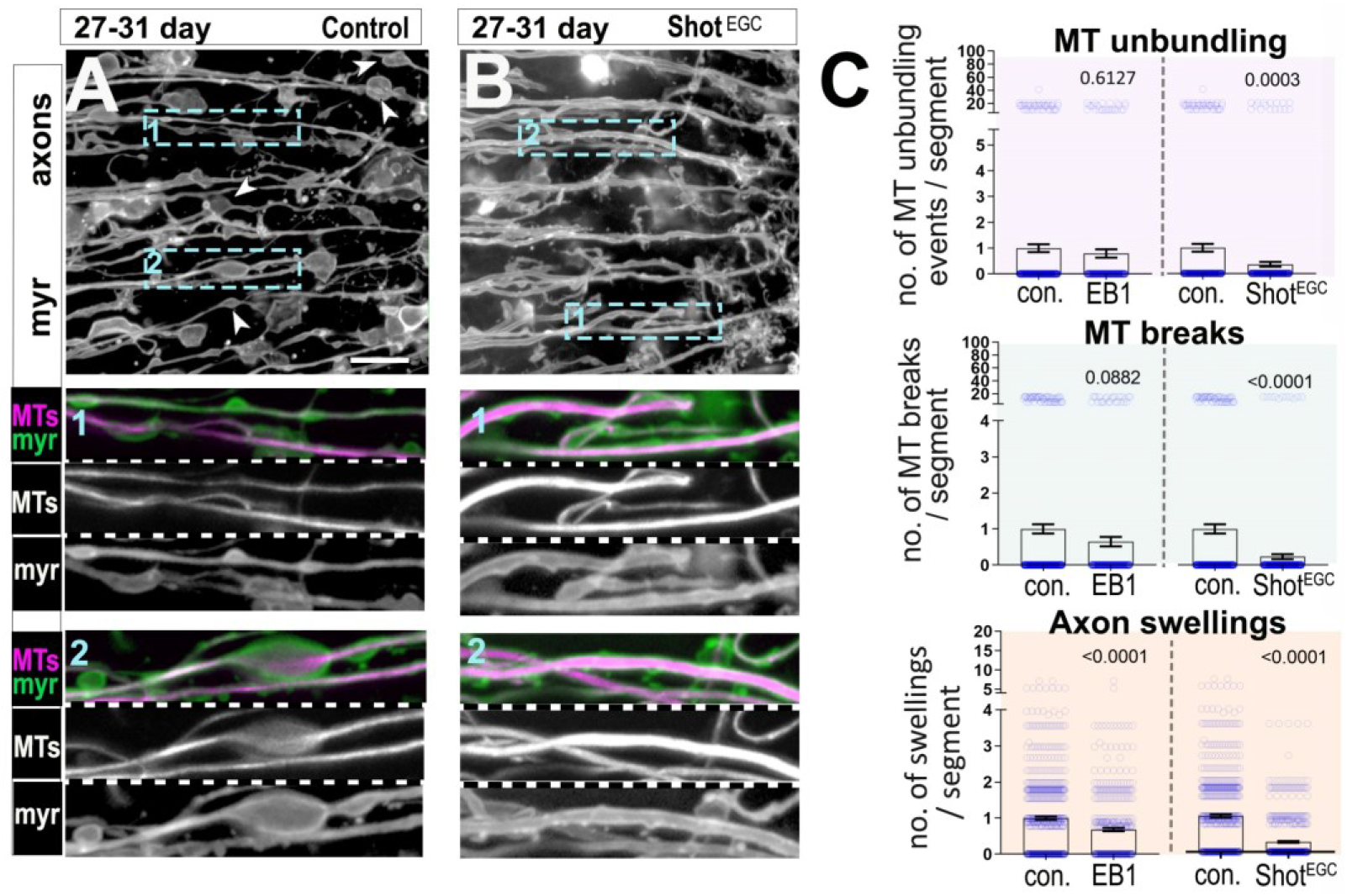
Post-developmental expression of Shot^EGC^ but not of EB1 is sufficient to rescue all axonal ageing phenotypes. (**A,B**) T1 axons in the medulla of old specimens of 27-33 days, are labelled with the plasma membrane marker myr-Tom (myr, greyscale images in A and B to visualise MTs, and green in double -channel images) and either GFP-tagged α-tubulin in A, or GFP-tagged Shot^EGC^ in B. Using the UAS/Gal4/Gal80^ts^ system, gene expression is induced after development by shifting newly hatched flies from 18 to 29°C. Ageing phenotypes including axon swellings, axon thinning (A), MT thinning (boxed area 1 in A) and MT unbundling (boxed area 2 in A) can be observed in old specimens, but are absent upon postdevelopment-expression of Shot^EGC^ in mature T1 neurons (A compared to B, boxed areas 1 and 2 shown as 2-fold magnified inset were MTs are labelled with GFP-tagged α-tubulin in A or GFP-tagged Shot^EGC^ in B). (**C**) Quantifications of phenotypes in old specimens shown in A and B, plus conditions of post-development expression of EB1 in old specimens (27-33 days old). Specific conditions are indicated below X-axes; Data points are shown as blue circles and as mean ± SEM; P-values obtained via Mann-Whitney test are indicated above. Data were taken from a minimum of 23 specimens per group. For detailed statistical values see table S13. Scale bar in A represents 10 μm.

