## Supplemental Tables for "Microtubule decay is a driver of neuronal ageing and a promising target for intervention"

| Genotype | *UAS-myr-tdTomato /+ ; UAS-GFP-α-tubulin84B, GMR31F10-Gal4/+* | | | |
|---|---|---|---|---|
| Age (days) | **5-10** | **19-23** | **29-31** | **38-42** |
| | **Axon diameter** | | | |
| Number of values (axons) | 145 | 143 | 104 | 241 |
| Number specimens | 10 | 14 | 11 | 29 |
| Mean | 0.7094 | 0.6716 | 0.4997 | 0.4878 |
| Std. Deviation | 0.1546 | 0.1 | 0.07939 | 0.1029 |
| Std. Error of Mean | 0.01284 | 0.008363 | 0.007785 | 0.00663 |
| | **Axon swellings** | | | |
| Number of values (μm of axon) | 303 | 303 | 240 | 319 |
| Number specimens | 24 | 22 | 24 | 28 |
| Mean | 0.007629 | 0.01310 | 0.01397 | 0.02427 |
| Std. Deviation | 0.01474 | 0.02056 | 0.02106 | 0.02773 |
| Std. Error of Mean | 0.0008466 | 0.001181 | 0.001359 | 0.001552 |
| | **Terminal area** | | | |
| Number of values (terminals) | 366 | 535 | 426 | 467 |
| Number specimens | 14 | 15 | 23 | 33 |
| Mean | 36.18 | 37.67 | 32.45 | 27.95 |
| Std. Deviation | 12.58 | 13.47 | 12.12 | 11.59 |
| Std. Error of Mean | 0.6575 | 0.5824 | 0.5874 | 0.5362 |
| | **Terminal morphology** | | | |
| Number of values (terminals) | 1238 | 826 | 1118 | 1696 |
| Number of normal | 826 | 814 | 1050 | 1501 |
| Number of swollen+broken | 32 | 12 | 68 | 195 |
| Number specimens | 14 | 15 | 23 | 31 |

**Table S1:** Correspondent statistical details for Fig. 1 "Axons and synaptic terminals within the *Drosophila* visual system deteriorate during ageing".

| Age (days) | **5** | **14** | **23** | **31** |
|---|---|---|---|---|
| Genotype | *UAS-GFP-α-tubulin84B /+ ; UAS- RedStinger, GMR31F10-Gal4/+* | | | |
| | **T1 neurons** | | | |
| Number of values-specimens | 12 | 5 | 10 | 11 |
| Mean | 140.8 | 230.2 | 200.6 | 241.0 |
| Std. Deviation | 39.38 | 40.55 | 41.04 | 38.26 |
| Std. Error of Mean | 11.37 | 18.14 | 12.98 | 11.54 |

**Table S2:** Correspondent statistical details for Fig. S2 "Absence of neuronal death amongst T1 neurons during ageing".

| Age (days) | **3 days 18$^0$C + 4 days 29$^0$C** | **56 days 18$^0$C + 4 days 29$^0$C** |
|---|---|---|
| Genotype | *tub-Gal80$^{ts}$/ UAS-myr-tdTomato; UAS-GFP-α-tubulin84B, GMR31F10-Gal4/+* | |
| | **Axon diameter** | |
| Number of values (axons) | 66 | 85 |
| Number specimens | 5 | 8 |
| Mean | 0.7732 | 0.5885 |
| Std. Deviation | 0.1393 | 0.1052 |
| Std. Error of Mean | 0.01714 | 0.01141 |
| | **Axon swellings** | |
| Number of values (μm of axon) | 123 | 98 |
| Number specimens | 5 | 8 |
| Mean | 0.006378 | 0.01722 |
| Std. Deviation | 0.01307 | 0.02129 |
| Std. Error of Mean | 0.001178 | 0.002151 |
| | **Terminal morphology** | |
| Number of values (terminals) | 300 | 485 |
| Number of normal | 298 | 371 |
| Number of swollen+broken | 2 | 114 |
| Number specimens | 5 | 8 |

**Table S3:** Correspondent statistical details for Fig. S3 "The deterioration of T1 neurons during ageing is independent of temperature and marker expression".

<table>
<tr><td>Age (days)</td><td>5-10</td><td>19-23</td><td>29-31</td><td>38-42</td></tr>
<tr><td>Genotype</td><td colspan="4">UAS-myr-tdTomato; UAS-GFP-α-tubulin84B, GMR31F10-Gal4/+</td></tr>
<tr><td></td><td colspan="4">MT unbundling</td></tr>
<tr><td>Number of values (axonal bundle segment)</td><td>562</td><td>713</td><td>616</td><td>264</td></tr>
<tr><td>Number specimens</td><td>22</td><td>23</td><td>17</td><td>17</td></tr>
<tr><td>Mean</td><td>0.08007</td><td>0.1445</td><td>0.1851</td><td>0.3030</td></tr>
<tr><td>Std. Deviation</td><td>0.3027</td><td>0.4404</td><td>0.4887</td><td>0.6034</td></tr>
<tr><td>Std. Error of Mean</td><td>0.01277</td><td>0.01649</td><td>0.01969</td><td>0.03714</td></tr>
<tr><td></td><td colspan="4">MT breaks</td></tr>
<tr><td>Number of values (axonal bundle segment)</td><td>563</td><td>696</td><td>500</td><td>266</td></tr>
<tr><td>Number specimens</td><td>22</td><td>23</td><td>17</td><td>17</td></tr>
<tr><td>Mean</td><td>0.04263</td><td>0.1710</td><td>0.1420</td><td>0.5489</td></tr>
<tr><td>Std. Deviation</td><td>0.2022</td><td>0.4562</td><td>0.4629</td><td>1.046</td></tr>
<tr><td>Std. Error of Mean</td><td>0.008522</td><td>0.01729</td><td>0.02070</td><td>0.06413</td></tr>
<tr><td></td><td colspan="4">Synaptic MTs</td></tr>
<tr><td>Number of values (terminal)</td><td>297</td><td>458</td><td>333</td><td>252</td></tr>
<tr><td>Number specimens</td><td>15</td><td>18</td><td>15</td><td>13</td></tr>
<tr><td>Mean</td><td>10.25</td><td>7.175</td><td>5.186</td><td>3.988</td></tr>
<tr><td>Std. Deviation</td><td>2.646</td><td>2.226</td><td>1.874</td><td>1.633</td></tr>
<tr><td></td><td colspan="4">MT bundle diameter Airyscan</td></tr>
<tr><td>Age (days)</td><td colspan="2">8-10</td><td colspan="2">34-36</td></tr>
<tr><td>Number of values (axons)</td><td colspan="2">57</td><td colspan="2">57</td></tr>
<tr><td>Number specimens</td><td colspan="2">14</td><td colspan="2">13</td></tr>
<tr><td>Mean</td><td colspan="2">0.7610</td><td colspan="2">0.6040</td></tr>
<tr><td>Std. Deviation</td><td colspan="2">0.1702</td><td colspan="2">0.1433</td></tr>
<tr><td>Std. Error of Mean</td><td colspan="2">0.02254</td><td colspan="2">0.01897</td></tr>
<tr><td></td><td colspan="4">MT bundle diameter STED</td></tr>
<tr><td>Age (days)</td><td colspan="2">3</td><td colspan="2">3</td></tr>
<tr><td>Number of values (axons)</td><td colspan="2">9</td><td colspan="2">8</td></tr>
<tr><td>Number specimens</td><td colspan="2">3</td><td colspan="2">3</td></tr>
<tr><td>Mean</td><td colspan="2">0.9265</td><td colspan="2">0.5552</td></tr>
<tr><td>Std. Deviation</td><td colspan="2">0.1785</td><td colspan="2">0.1255</td></tr>
<tr><td>Std. Error of Mean</td><td colspan="2">0.05950</td><td colspan="2">0.04438</td></tr>
</table>

**Table S4:** Correspondent statistical details for Fig. 2 “MT aberration precedes axonal and synaptic decay during ageing”

| Age | **3 days 8$^0$C + 15 days 29$^0$C** | **56 days 18$^0$C + 15 days 29$^0$C** |
|---|---|---|
| Genotype | *tub-Gal80$^{ts}$/ UAS-myr-tdTomato*; *UAS-GFP-α-tubulin84B, GMR31F10-Gal4/+* | |
| | **MT unbundling** | |
| Number of values (axonal bundle segment) | 108 | 132 |
| Number specimens | 6 | 8 |
| Mean | 0.04630 | 0.3636 |
| Std. Deviation | 0.2515 | 0.6910 |
| Std. Error of Mean | 0.02420 | 0.06014 |
| | **MT breaks** | |
| Number of values (axonal bundle segment) | 108 | 132 |
| Number specimens | 6 | 8 |
| Mean | 0.009259 | 0.3864 |
| Std. Deviation | 0.09623 | 0.9050 |
| Std. Error of Mean | 0.009259 | 0.07877 |
| | **Synaptic MTs** | |
| Number of values (terminal) | 108 | 132 |
| Number specimens | 6 | 8 |
| Mean | 7.350 | 3.994 |
| Std. Deviation | 2.285 | 1.643 |
| Std. Error of Mean | 0.1931 | 0.09044 |

**Table S5:** Correspondent statistical details for Fig. S5. "MT alterations during ageing are independent of the MT reporter and are specific for aged specimens".

| Age (days) | **29-35** | |
|---|---|---|
| Genotype | *UAS-myr-tdTomato /+ ; UAS-GFP-α-tubulin84B, GMR31F10-Gal4/+* | *UAS-myr-tdTomato /+ ; UAS-GFP-α-tubulin84B, GMR31F10-Gal4/* ***UAS-mth$^{RNAi}$*** |
| | **Axon swellings** | |
| Number of values (axonal bundle segment) | 520 | 455 |
| Number specimens | 23 | 16 |
| Mean | 1.000 | 0.6137 |
| Std. Deviation | 1.531 | 1.092 |
| Std. Error of Mean | 0.06714 | 0.05121 |
| | **MT breaks** | |
| Number of values (axonal bundle segment) | 535 | 459 |
| Number specimens | 23 | 16 |

| | | |
|---|---|---|
| Mean | 1.000 | 0.2565 |
| Std. Deviation | 3.562 | 1.080 |
| Std. Error of Mean | 0.1540 | 0.05042 |
| | **MT unbundling** | |
| Number of values (axonal bundle segment) | 523 | 460 |
| Number specimens | 23 | 16 |
| Mean | 0.9999 | 0.6256 |
| Std. Deviation | 2.758 | 1.860 |
| Std. Error of Mean | 0.1206 | 0.08674 |
| | **Axon diameter** | |
| Age (days) | **33-41** | |
| Number of values (segment) | 137 | 116 |
| Number specimens | 12 | 10 |
| Mean | 1.000 | 1.323 |
| Std. Deviation | 0.2150 | 0.2626 |
| Std. Error of Mean | 0.01837 | 0.02438 |
| | **Synaptic MTs** | |
| Age (days) | **33-35** | |
| Number of values (terminal) | 231 | 211 |
| Number specimens | 11 | 10 |
| Mean | 4.719 | 7.479 |
| Std. Deviation | 1.773 | 2.183 |
| Std. Error of Mean | 0.1166 | 0.1503 |

**Table S6:** Correspondent statistical details for Fig. 3 "Knockdown of the ageing gene *mth* improves neuronal ageing hallmarks and MT decay".

| | Ratio GFP/Tom | |
|---|---|---|
| Age | **3 days 18$^0$C + 4 days 29$^0$C** | **56 days 18$^0$C + 4 days 29$^0$C** |
| Genotype | *tub-Gal80$^{ts}$/ UAS-myr-tdTomato; UAS-GFP-α-tubulin84B, GMR31F10-Gal4/+* | |
| Number of values (axons)<br>Number specimens<br>Mean<br>Std. Deviation<br>Std. Error of Mean | 127<br>5<br>1.848<br>0.4595<br>0.04078 | 200<br>7<br>1.076<br>0.2338<br>0.01653 |
| | **Ratio GFP/Tom** | |
| Age | **3 days 18$^0$C + 15 days 29$^0$C** | **56 days 18$^0$C + 15 days 29$^0$C** |
| Genotype | *tub-Gal80$^{ts}$/ UAS-myr-tdTomato; UAS-GFP-α-tubulin84B, GMR31F10-Gal4/+* | |
| Number of values (axons)<br>Number specimens<br>Mean<br>Std. Deviation<br>Std. Error of Mean | 126<br>6<br>1.701<br>0.5551<br>0.04946 | 225<br>8<br>1.352<br>0.4969<br>0.03313 |

**Table S7:** Correspondent statistical details for Fig. 4 “Decreased presence of tubulin-GFP at axonal microtubules in a pulse chase experiment suggests changes in MT turnover with age”.

| | EB1 in axons | |
|---|---|---|
| Age | **3-6 days** | **28-35 days** |
| Genotype | ***WT*** | |
| Number of values (medullas)<br>Number specimens<br>Mean<br>Std. Deviation<br>Std. Error of Mean | 26<br>15<br>1.000<br>0.2688<br>0.05271 | 29<br>17<br>0.8534<br>0.3349<br>0.06219 |
| | **Tau in axons** | |
| Age | **13-17 days 25$^0$C** | **75-83 days 25$^0$C** |
| Genotype | **Tau$^{wee304}$/+** | |
| Number of values (medullas)<br>Number specimens<br>Mean<br>Std. Deviation<br>Std. Error of Mean | 31<br>17<br>45.57<br>45.34<br>8.143 | 29<br>15<br>21.46<br>35.92<br>6.670 |
| | **Total EB1 (WB)** | |
| Age | **3-9 days** | **28-34 days** |
| Genotype | ***WT*** | |
| Number of values<br>Mean<br>Std. Deviation<br>Std. Error of Mean | 6<br>1.024<br>0.1688<br>0.06890 | 6<br>1.080<br>0.4631<br>0.1891 |

| | Total Tau (WB) | |
|---|---|---|
| Age | **6-9 days** | **30-35 days** |
| Genotype | ***WT*** | |
| Number of values<br>Mean<br>Std. Deviation<br>Std. Error of Mean | 10<br>0.9664<br>0.1582<br>0.05002 | 11<br>0.6505<br>0.2655<br>0.08005 |
| | **EB1-MT enrichment assay** | |
| Age | **2-6 days** | **28-31 days** |
| Genotype | ***WT*** | |
| Number of values<br>**Mean MT pellet**<br>Std. Deviation<br>Std. Error of Mean | 9<br>1.110<br>0.3633<br>0.1211 | 10<br>0.5600<br>0.2073<br>0.06556 |
| **Mean supernatant** | 1.062 | 1.069 |
| Std. Deviation | 0.1577 | 0.2732 |
| Std. Error of Mean | 0.04988 | 0.08638 |
| | **Tau-MT enrichment assay** | |
| Age | **2-6 days** | **28-31 days** |
| Genotype | ***WT*** | |
| Number of values<br>**Mean MT pellet**<br>Std. Deviation<br>Std. Error of Mean | 10<br>1.193<br>0.3165<br>0.1001 | 10<br>0.8550<br>0.2171<br>0.06866 |
| **Mean supernatant** | 1.062 | 1.069 |
| Std. Deviation | 0.1577 | 0.2732 |
| Std. Error of Mean | 0.04988 | 0.08638 |

**Table S8:** Correspondent statistical details for Fig. 5 “The function of EB1 and Tau is altered during ageing”.

| Genotype | *UAS-GFP-α-tubulin84B, GMR31F10-Gal4/+* | *UAS-GFP-α-tubulin84B, GMR31F10-Gal4/UAS-**tau**$^{RNAi}$* | *UAS-myr-tdTomato /+ ; UAS-GFP-α-tubulin84B, GMR31F10-Gal4/+* | *UAS-myr-tdTomato /UAS-**shot**$^{RNAi}$ ; UAS-GFP-α-tubulin84B, GMR31F10-Gal4/+* |
|---|---|---|---|---|
| Age (days) | **30-37** | **30-37** | **30-33** | **30-33** |
| | **Axon swellings** | | | |
| Number of values (axonal bundle segment) | 511 | 502 | 310 | 295 |
| Number specimens | 19 | 17 | 14 | 12 |
| Mean | 1.000 | 1.278 | 1.001 | 2.109 |
| Std. Deviation | 1.198 | 1.389 | 1.440 | 2.566 |
| Std. Error of Mean | 0.05302 | 0.06198 | 0.08179 | 0.1494 |
| | **MT unbundling** | | | |
| Number of values (axonal bundle segment) | 511 | 502 | 310 | 289 |
| Number specimens | 19 | 17 | 14 | 12 |
| Mean | 1.000 | 3.885 | 0.9999 | 4.704 |
| Std. Deviation | 5.597 | 12.42 | 2.810 | 8.794 |
| Std. Error of Mean | 0.2476 | 0.5542 | 0.1596 | 0.5173 |
| | **MT breaks** | | | |
| Number of values (axonal bundle segment) | 511 | 502 | 311 | 289 |
| Number specimens | 19 | 17 | 14 | 12 |
| Mean | 1.000 | 1.421 | 0.9967 | 1.756 |
| Std. Deviation | 3.280 | 3.686 | 2.720 | 3.848 |
| Std. Error of Mean | 0.1451 | 0.1645 | 0.1543 | 0.2229 |

| Genotype | *UAS-myr-tdTomato /+ ; UAS-GFP-α-tubulin84B, GMR31F10-Gal4/+* | *UAS-myr-tdTomato / UAS-**shot**$^{RNAi}$ ; UAS-GFP-α-tubulin84B, GMR31F10-Gal4/UAS-**Tau**$^{RNAi}$* | *UAS-myr-tdTomato /+ ; UAS-GFP-α-tubulin84B, GMR31F10-Gal4/+* | *UAS-myr-tdTomato **/EB1 Df**; UAS-GFP-α-tubulin84B, GMR31F10-Gal4/ **UAS-EB1**$^{RNAi}$* |
|---|---|---|---|---|
| Age (days) | **28-32** | **28-32** | **25-28** | **25-28** |
| | **Axon swellings** | | | |
| Number of values (axonal bundle segment) | 340 | 446 | 464 | 457 |
| Number specimens | 19 | 18 | 14 | 14 |
| Mean | 1.000 | 3.366 | 1.000 | 2.907 |
| Std. Deviation | 1.806 | 3.892 | 2.351 | 4.458 |

| | | | | |
|---|---|---|---|---|
| Std. Error of Mean | 0.09792 | 0.1843 | 0.1091 | 0.2085 |
| | **MT unbundling** | | | |
| Number of values (axonal bundle segment) | 340 | 442 | 465 | 461 |
| Number specimens | 19 | 18 | 14 | 14 |
| Mean | 1.000 | 5.103 | 1.000 | 2.143 |
| Std. Deviation | 3.237 | 8.556 | 3.639 | 6.860 |
| Std. Error of Mean | 0.1756 | 0.4070 | 0.1687 | 0.3195 |
| | **MT breaks** | | | |
| Number of values (axonal bundle segment) | 344 | 437 | 466 | 465 |
| Number specimens | 19 | 18 | 14 | 14 |
| Mean | 0.9999 | 3.173 | 1.000 | 2.795 |
| Std. Deviation | 3.013 | 6.000 | 3.267 | 5.571 |
| Std. Error of Mean | 0.1622 | 0.2870 | 0.1514 | 0.2583 |

**Table S9:** Correspondent statistical details for Fig. 6 “Reduction of Eb1, Tau and Shot exacerbates age-related MT decay and ageing hallmarks”.

| Genotype | *UAS-myr-tdTomato /+ ; UAS-GFP-α-tubulin84B, GMR31F10-Gal4/+* | *UAS-myr-tdTomato / UAS-**Shot**$^{RNi}$ ; UAS-GFP-α-tubulin84B, GMR31F10-Gal4/UAS-**tau**$^{RNAi}$* | *UAS-myr-tdTomato /+ ; UAS-GFP-α-tubulin84B, GMR31F10-Gal4/+* | *UAS-myr-tdTomato /+; UAS-GFP-α-tubulin84B, GMR31F10-Gal4/ **UAS-mcherry-EB1*** |
|---|---|---|---|---|
| Age (days) | **28-32** | **28-32** | **25-28** | **25-28** |
| | **Synaptic MTs** | | | |
| Number of values (terminals) | 340 | 446 | 219 | 184 |
| Number specimens | 14 | 20 | 17 | 14 |
| Mean | 5.703 | 2.574 | 4.909 | 6.658 |
| Std. Deviation | 2.116 | 1.598 | 1.668 | 1.761 |
| Std. Error of Mean | 0.1614 | 0.1027 | 0.1127 | 0.1298 |
| | **Terminal Morphology** | | | |
| Number of values (terminals) | 1143 | 1220 | 2006 | 1770 |
| Number of normal | 1028 | 663 | 1506 | 1623 |
| Number of swollen+broken | 115 | 557 | 500 | 147 |
| Number specimens | 14 | 20 | 17 | 14 |

| Genotype | *UAS-myr-tdTomato /+ ; UAS-GFP-α-tubulin84B, GMR31F10-Gal4/+* | *UAS-myr-tdTomato / +; GMR31F10-Gal4/ **UAS-Shot**$^{EGC}$**-GFP*** |
|---|---|---|
| Age (days) | **27-31** | **27-31** |
| | **Synaptic MTs** | |
| Number of values (terminals) | 194 | 146 |
| Number specimens | 15 | 13 |
| Mean | 5.701 | 6.411 |
| Std. Deviation | 2.057 | 2.237 |
| Std. Error of Mean | 0.1477 | 0.1851 |
| | **Terminal Morphology** | |
| Number of values (terminals) | 340 | 442 |
| Number of normal | 725 | 654 |
| Number of swollen+broken | 188 | 78 |
| Number specimens | 15 | 13 |

**Table S10:** Correspondent statistical details for Fig. S6 "The deterioration of synaptic terminals during ageing can be exacerbated or rescued by altering the function of MT regulators"

| Genotype | *UAS-GFP-α-tubulin84B, GMR31F10-Gal4/+* | *UAS-GFP-α-tubulin84B, GMR31F10-Gal4/UAS-**tau**$^{RNAi}$* | *UAS-myr-tdTomato /+ ; UAS-GFP-α-tubulin84B, GMR31F10-Gal4/+* | *UAS-myr-tdTomato /UAS-s**hot**$^{RNAi}$ ; UAS-GFP-α-tubulin84B, GMR31F10-Gal4/+* |
|---|---|---|---|---|
| Age (days) | **3-8** | **3-8** | **5-7** | **5-7** |
| | **Axon swellings** | | | |
| Number of values (axonal bundle segment) | 567 | 598 | 73 | 311 |
| Number specimens | 19 | 24 | 6 | 8 |
| Mean | 1.000 | 1.102 | 0.9999 | 1.056 |
| Std. Deviation | 1.766 | 2.280 | 1.948 | 2.154 |
| Std. Error of Mean | 0.07417 | 0.09325 | 0.2280 | 0.1221 |
| | **MT unbundling** | | | |
| Number of values (axonal bundle segment) | 478 | 598 | 73 | 319 |
| Number specimens | 19 | 24 | 6 | 8 |
| Mean | 1.000 | 0.1802 | 1.000 | 0.5035 |
| Std. Deviation | 4.789 | 1.963 | 3.714 | 2.668 |
| Std. Error of Mean | 0.2191 | 0.08029 | 0.4346 | 0.1494 |
| | **MT breaks** | | | |
| Number of values (axonal bundle segment) | 478 | 597 | 73 | 321 |
| Number specimens | 19 | 24 | 6 | 8 |
| Mean | 1.000 | 1.021 | 0.9999 | 0.6822 |
| Std. Deviation | 2.054 | 2.118 | 4.863 | 4.023 |
| Std. Error of Mean | 0.09393 | 0.08668 | 0.5692 | 0.2245 |
| Genotype | *UAS-myr-tdTomato /+ ; UAS-GFP-α-tubulin84B, GMR31F10-Gal4/+* | *UAS-myr-tdTomato / UAS-**Shot**$^{RNI}$ ; UAS-GFP-α-tubulin84B, GMR31F10-Gal4/UAS-**tau**$^{RNAi}$* | *UAS-myr-tdTomato /+ ; UAS-GFP-α-tubulin84B, GMR31F10-Gal4/+* | *UAS-myr-tdTomato **/EB1 Df**; UAS-GFP-α-tubulin84B, GMR31F10-Gal4/ **UAS-eb1**$^{RNAi}$* |
| Age (days) | **3-7** | **3-7** | **3-8** | **3-8** |
| | **Axon swellings** | | | |
| Number of values (axonal bundle segment) | 383 | 450 | 329 | 359 |
| Number specimens | 17 | 18 | 13 | 17 |
| Mean | 0.9973 | 0.7563 | 1.000 | 0.7590 |
| Std. Deviation | 1.065 | 1.080 | 1.366 | 1.176 |

| | | | | |
|---|---|---|---|---|
| Std. Error of Mean | 0.05449 | 0.05093 | 0.07533 | 0.06234 |
| | **MT unbundling** | | | |
| Number of values (axonal bundle segment) | 382 | 450 | 329 | 357 |
| Number specimens | 17 | 18 | 13 | 17 |
| Mean | 1.000 | 1.313 | 0.9999 | 0.4280 |
| Std. Deviation | 8.320 | 9.268 | 5.362 | 4.219 |
| Std. Error of Mean | 0.4257 | 0.4369 | 0.2956 | 0.2233 |
| | **MT breaks** | | | |
| Number of values (axonal bundle segment) | 382 | 450 | 329 | 356 |
| Number specimens | 17 | 18 | 13 | 17 |
| Mean | 1.000 | 1.352 | 1.000 | 0.7732 |
| Std. Deviation | 8.504 | 10.80 | 2.071 | 1.837 |
| Std. Error of Mean | 0.4351 | 0.5093 | 0.1142 | 0.09734 |

**Table S11:** Correspondent statistical details for Fig. S7 “MT decay and ageing hallmarks are not present in young specimens with EB1, Tau and Shot knockdowns”.

| Genotype | *UAS-myr-tdTomato /+ ; UAS-GFP-α-tubulin84B, GMR31F10-Gal4/+* | *UAS-myr-tdTomato /+; UAS-GFP-α-tubulin84B, GMR31F10-Gal4/* ***UAS-mcherry-EB1*** | *UAS-myr-tdTomato /+ ; UAS-GFP-α-tubulin84B, GMR31F10-Gal4/+* | *UAS-myr-tdTomato / +; GMR31F10-Gal4/* ***UAS-Shot$^{EGC}$-GFP*** |
|---|---|---|---|---|
| Age (days) | **28-30** | **28-30** | **27-31** | **27-31** |
| | **MT unbundling** | | | |
| Number of values (axonal bundle segment) | 377 | 308 | 382 | 303 |
| Number specimens | 17 | 14 | 14 | 13 |
| Mean | 0.9999 | 0.4584 | 1.000 | 0.03287 |
| Std. Deviation | 2.579 | 1.938 | 2.825 | 0.4039 |
| Std. Error of Mean | 0.1328 | 0.1104 | 0.1445 | 0.02321 |
| | **MT breaks** | | | |
| Number of values | 371 | 302 | 238 | 241 |
| Number specimens | 17 | 14 | 14 | 13 |
| Mean | 0.9421 | 0.4247 | 0.9998 | 0.4481 |
| Std. Deviation | 2.996 | 1.779 | 2.844 | 1.744 |
| Std. Error of Mean | 0.1555 | 0.1024 | 0.1844 | 0.1123 |
| | **Axon Swellings** | | | |
| Number of values (axonal bundle segment) | 375 | 301 | 381 | 303 |

| | | | | |
|---|---|---|---|---|
| Number specimens | 17 | 14 | 14 | 13 |
| Mean | 0.9999 | 0.6126 | 1.000 | 0.01488 |
| Std. Deviation | 1.499 | 1.076 | 1.828 | 0.1829 |
| Std. Error of Mean | 0.07742 | 0.06201 | 0.09366 | 0.01051 |
| | **MT bundle diameter** | | | |
| Age (days) | **25-28** | **25-28** | **29-30** | **29-30** |
| Number of values (axonal bundle segment) | 31 | 40 | 145 | 129 |
| Number specimens | 5 | 7 | 10 | 10 |
| Mean | 1.000 | 1.261 | 1.000 | 1.377 |
| Std. Deviation | 0.2152 | 0.3327 | 0.2281 | 0.2584 |
| Std. Error of Mean | 0.03865 | 0.05261 | 0.01894 | 0.02275 |
| | **Axon diameter** | | | |
| Age (days) | **35-38** | **35-38** | **27-31** | **27-31** |
| Number of values (axons) | 291 | 165 | 207 | 242 |
| Number specimens | 16 | 12 | 13 | 12 |
| Mean | 1.000 | 1.137 | 1.000 | 1.468 |
| Std. Deviation | 0.1673 | 0.1976 | 0.1789 | 0.2858 |
| Std. Error of Mean | 0.009805 | 0.01539 | 0.01243 | 0.01837 |

**Table S12:** Correspondent statistical details for Fig. 7 "Expression of EB1 and Shot$^{EGC}$ ameliorates axonal ageing phenotypes"

| Genotype | *tub-Gal80^ts/ UAS-myr-tdTomato; UAS-GFP-α-tubulin84B, GMR31F10-Gal4/+* | *tub-Gal80^ts/ UAS-myr-tdTomato; UAS-GFP-α-tubulin84B, GMR31F10-Gal4/* ***UAS-mcherry-EB1*** | *tub-Gal80^ts/ UAS-myr-tdTomato; UAS-GFP-α-tubulin84B, GMR31F10-Gal4/+* | *tub-Gal80^ts/ UAS-myr-tdTomato; GMR31F10-Gal4/* ***UAS-Shot^EGC-GFP*** |
|---|---|---|---|---|
| Age (days) | **27-31** | **27-31** | **27-31** | **27-31** |
| | **MT unbundling** | | | |
| Number of values (axonal bundle segment) | 721 | 449 | 721 | 697 |
| Number specimens | 34 | 24 | 26 | 23 |
| Mean | 1.000 | 0.7952 | 1.000 | 0.3564 |
| Std. Deviation | 4.104 | 3.441 | 4.104 | 2.440 |
| Std. Error of Mean | 0.1529 | 0.1624 | 0.1529 | 0.09242 |
| | **MT breaks** | | | |
| Number of values (axonal bundle segment) | 721 | 449 | 721 | 697 |
| Number specimens | 34 | 24 | 26 | 23 |
| Mean | 1.000 | 0.6500 | 1.000 | 0.2334 |
| Std. Deviation | 3.441 | 2.785 | 3.441 | 1.849 |
| Std. Error of Mean | 0.1282 | 0.1314 | 0.1282 | 0.07003 |
| | **Axon Swellings** | | | |
| Number of values (axonal bundle segment) | 721 | 449 | 721 | 697 |
| Number specimens | 34 | 24 | 26 | 23 |
| Mean | 1.000 | 0.6801 | 1.000 | 0.2759 |
| Std. Deviation | 1.148 | 0.9823 | 1.148 | 0.6025 |
| Std. Error of Mean | 0.04274 | 0.04636 | 0.04274 | 0.02282 |

**Table S13:** Correspondent statistical details for Fig. S8 "Post-developmental expression of Shot$^{EGC}$ but not of EB1 is sufficient to rescue all axonal ageing phenotypes".

| Genotype | *tub-Gal80$^{ts}$/ +; elav-Gal4/+* | *tub-Gal80$^{ts}$/ +; elav-Gal4/**UAS-mcherry-EB1*** | *tub-Gal80$^{ts}$/ +; elav-Gal4/**UAS-Shot$^{EGC}$-GFP*** | *tub-Gal80$^{ts}$/ +; elav-Gal4/+* | *tub-Gal80$^{ts}$/ +; elav-Gal4/**UAS-mcherry-EB1*** | *tub-Gal80$^{ts}$/ +; elav-Gal4/**UAS-Shot$^{EGC}$-GFP*** |
|---|---|---|---|---|---|---|
| Age (days) | **4-5** | **4-5** | **4-5** | **25-26** | **25-26** | **25-26** |
| Number specimens | 239 | 162 | 146 | 230 | 143 | 168 |
| Mean | 15.00 | 10.02 | 14.71 | 1.004 | 0.9091 | 2.012 |
| Std. Deviation | 4.733 | 5.817 | 4.489 | 1.884 | 1.524 | 2.189 |
| Std. Error of Mean | 0.3061 | 0.4570 | 0.3715 | 0.1242 | 0.1274 | 0.1689 |

**Table S14:** Correspondent statistical details for Fig. 8 "The decline in locomotion of aged flies improves upon adult expression of Shot$^{EGC}$ ".
